# Modulating the Kynurenine pathway or sequestering toxic 3-hydroxykynurenine protects the retina from light induced damage in *Drosophila*

**DOI:** 10.1101/2022.09.10.507411

**Authors:** Sarita Hebbar, Sofia Traikov, Catrin Hälsig, Elisabeth Knust

## Abstract

Tissue health is regulated by a myriad of exogenous or endogenous factors. Here we investigated the role of the conserved Kynurenine pathway (KP) in maintaining retinal homeostasis in the context of light stress in *Drosophila melanogaster*. *cinnabar, cardinal* and, *scarlet*, are fly genes that encode different steps in the KP. Along with *white*, these genes are known regulators of brown pigment (ommochrome) biosynthesis. Using *white* as a sensitized genetic background, we showed that mutations in *cinnabar, cardinal*, and *scarlet* differentially modulate light-induced retinal damage. Mass Spectrometric measurements of KP metabolites in flies with different genetic combinations support the notion that increased levels of 3-hydroxykynurenine (3OH-K) and Xanthurenic acid (XA) enhance retinal damage, whereas Kynurenic Acid (KYNA) and Kynurenine (K) are neuro-protective. This conclusion was corroborated by showing that feeding 3OH-K results in enhanced retinal damage, whereas feeding KYNA protects the retina in sensitized genetic backgrounds. Interestingly, the harmful effects of free 3OH-K are diminished by its sub-cellular compartmentalization within the cell. Sequestering of 3OH-K enables the quenching of its toxicity through conversion to brown pigment or conjugation to proteins. This work enabled us to decouple the role of these KP genes in ommochrome formation from their role in retinal homeostasis. Additionally, it puts forward new hypotheses on the importance of the balance of KP metabolites and their compartmentalization in disease alleviation.

## Introduction

Tissue health is regulated by a myriad of exogenous and endogenous factors, and deregulation of signaling and metabolic pathways have been linked to human diseases. There is increasing evidence that metabolic pathways play crucial roles in the progression of, or protection from, neurodegenerative diseases, including Parkinson’s and Alzheimer’s disease (Camandola & Mattson, 2017) or retinal diseases (Fiedorowicz, Choragiewicz et al., 2019, Li, Cai et al., 2021, Pan, Wubben et al., 2021, Wubben, Pawar et al., 2017). In particular, the Kynurenine pathway (KP) has more recently attracted much attention. It is one of the pathways downstream of tryptophan (Trp), and its dysregulation can lead to accumulation of intermediates, which can be either toxic or beneficial and thus can exacerbate or ameliorate the phenotypic outcome of neurodegeneration (Davis & Liu, 2015). In short, Trp is converted by tryptophan 2,3 dioxygenase to N-formyl-L-kynurenine, which is, in a second step, hydrolyzed to kynurenine (K) by kynurenine formamidase (Jakoby, 1954), followed by conversion to 3-hydroxykynurenine (3OH-K), catalyzed by kynurenine 3-monooxygenase (KMO). Kynurenic Acid (KYNA) and Xanthurenic acid (XA) are other metabolites in this pathway, which can be produced from K and 3OH-K (Umebachi, 1985), respectively (Fig. 1A). K and 3OH-K have been associated with degenerative conditions in various animal models and in human diseased tissues (Colin-Gonzalez, Maldonado et al., 2013). The KP is under consideration as a therapeutic target in neurological disorders (Dantzer, 2017, Stone & Darlington, 2013). A major conundrum, however, is how these metabolites, for example K or 3OH-K, play modulating, sometimes opposing roles on the disease phenotype (Behl, Kaur et al., 2021, Boros, Bohar et al., 2018, Huang, Ogbechi et al., 2020, Mithaiwala, Santana-Coelho et al., 2021, Mor, Tankiewicz-Kwedlo et al., 2021). Other relevant questions in this context include: How do external factors, such as age or environmental stress, intersect in KP modulation of tissue health? What are the underlying cellular conditions for 3OH-K to exert its toxicity? What are the intracellular interactions of 3OH-K with biomolecules? How do these interactions regulate tissue health?

**Figure 1:**
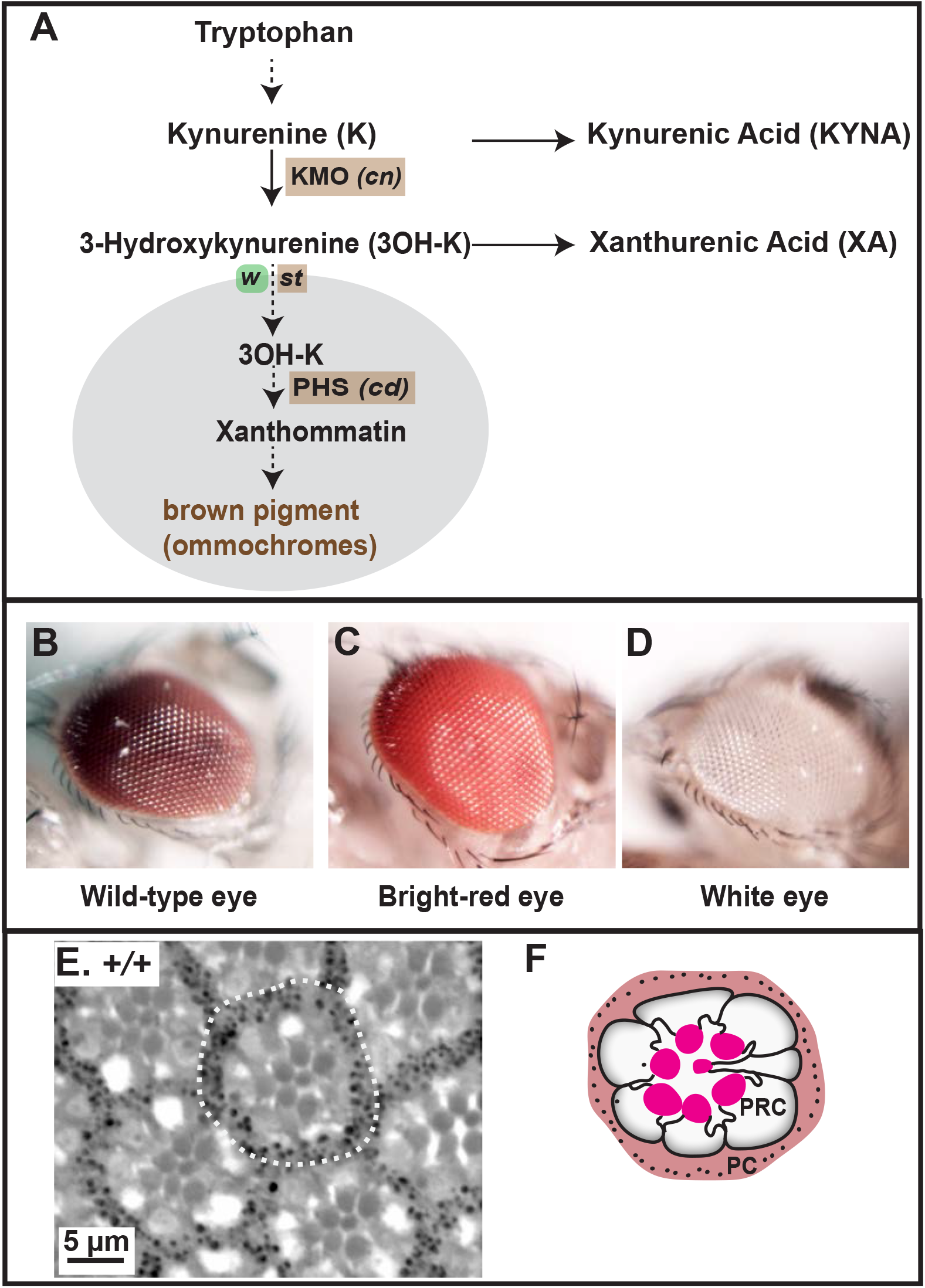
The Kynurenine pathway (KP) pathway in the context of brown pigment biogenesis in the fly eye. **A:** Relevant steps of the KP [from the KEGG database (Kanehisa, Sato et al., 2016)], starting from Tryptophan and leading to brown pigments/ommochromes. Solid arrows: direct steps as precursor-product conversion; dotted arrows: formation of a relevant product after a series of steps. Kynurenine 3-monooxygenase (KMO), and Phenoxazinone synthetase (PHS) are enzymes encoded by the *Drosophila* genes *cinnabar* (*cn*) and *cardinal* (*cd*), respectively. White and Scarlet, encoded by *white* (*w*) and *scarlet* (*st*), respectively, compartmentalize free 3OH-K through their transporter activity, allowing the formation of the brown pigment (ommochromes) within lysosomal-related organelles (LRO; grey). **B-D:** Photographs depicting eye color phenotypes of the mutants used in this study. (B) Wild type eye color, (C) bright-red eye color observed in *st* and *cd* alleles when brown pigment biogenesis is disrupted, (D) non-pigmented, white eye observed when brown and red pigment biogenesis is disrupted as in mutant alleles of *white*. **E-F**: Bright-field image (E) of a toluidine blue-stained 1μm thick section of a wild-type adult eye after exposure to continuous, high intensity light, showing several ommatidia. One ommatidium is marked with a dotted outline. Note the seven rhabdomeres in the center. (F) Corresponding schematic of the section in E, showing one ommatidium with seven photoreceptor cells (PRC) with their rhabdomeres (bright red), surrounded by pigment cells (PC).

Here, we address some of the above questions using *Drosophila melanogaster* and thus extend previous work done on KP. In *Drosophila melanogaster*, studies on the KP date back to the 1940s when it was shown that mutations in the eye color gene *vermilion* (*v*), which turns the wild-type eye color from dark red (Fig. 1B) to bright red (similar to that shown in Fig. 1C), affect the conversion of Trp to Kynurenine (K) (Ephrussi, 1942) (Fig. 1A). It was later shown that *v* encodes tryptophan 2,3 dioxygenase (Glassman, 1956). Beside *v*, several other eye color mutants resulting in bright red eyes turned out to be part of the KP pathway also: *cinnabar* (*cn*) encodes kynurenine 3-monooxygenase (KMO), which converts K to 3-hydroxykynurenine (3OH-K), *scarlet* (*st*) encodes a transporter of 3OH-K, and *cardinal* (*cd*) encodes a Phenoxazinone synthetase (PHS) (Fig. 1A). The eye color phenotype of *v* flies as well flies carrying mutations in *cn* or *st* (light red instead of dark red) is caused by the lack of the brown pigments, the ommochromes, which, together with the red pigments, the drosopterins, form the screening pigments in the *Drosophila* eye (Shoup, 1966). Screening pigments accumulate in lysosome-related organelles (LROs) within pigment cells, support cells that surround the centrally located photoreceptor cells (PRCs; Fig. 1E, F). These pigments shield PRCs from high intensity light. Formation of the brown pigment called xanthommatin in LROs depends on the transport of 3OH-K into LROs. This is mediated by proteins encoded by *scarlet* (*st*) and *white* (*w*) genes, orthologues of the heterodimeric ABC (ATP-binding cassette) transporters (Mackenzie, Brooker et al., 1999, Pepling & Mount, 1990, Tearle, Belote et al., 1989) (Fig. 1A). The subsequent action of Phenoxazinone synthetase, encoded by *cardinal* (*cd*), results in the formation of xanthommatin (Phillips, Forrest et al., 1973) which eventually gets converted to the brown pigment (ommochromes).

Strikingly, beside their role in ommochrome biosynthesis, *Drosophila cn*/KMO and *st* can modulate degeneration in neurons in disease models such as Huntington’s (Campesan, Green et al., 2011, Green, Campesan et al., 2012) and Parkinson’s Disease (Cunningham, Waldeck et al., 2018). These data together with the high degree of genetic conservation of disease genes in *Drosophila* support the fly model as a valuable system to unravel the underlying genetic and molecular basis of neurodegenerative diseases (Nitta & Sugie, 2022, Ugur, Chen et al., 2016), including retinal degeneration (Lehmann, Knust et al., 2019).

Both in human and in flies the severity of a mutant phenotype/symptoms in individuals carrying the same mutation can be highly variable, ranging from complete neuronal degeneration to weaker manifestations of the disease phenotype. This could be traced back to the genetic background, which can modulate the severity of a mutant phenotype in flies (Chow, Kelsey et al., 2016), mice (Schon, Asteriti et al., 2016) and human (Venturini, Rose et al., 2012). For example, mutations in *Drosophila w*, which lack all pigments and make the eyes white in color (Fig. 1D), modulate retinal degeneration in flies expressing the human Tau protein (Ambegaokar & Jackson, 2010). This suggests that mutations in *w* sensitize the eyes for degeneration. Given that the KP not only participates in brown pigment formation, but is also part of metabolic processes in cells, we hypothesize that the role of *w* in predisposing eyes to degeneration is linked to an imbalance in some KP metabolites, rather than to its role in pigment biosynthesis and hence shielding the eye from excess light. Data presented here using genetic interactions, biochemical analysis of metabolites and dietary intervention support this hypothesis. These approaches enabled us to identify a correlation between higher abundances of 3-hydroxy Kynurenine (3OH-K) under physiological conditions and the severity of retinal damage upon light stress conditions. The causal relationship between metabolite abundance and phenotype could be further documented by showing that dietary supplementation of 3OH-K modulates the degree of light-induced damage. Our results further suggest that free 3OH-K, rather than compartmentalized and protein-bound 3OH-K, is particularly detrimental to the retina.

## Results

### The degree of light-induced damage in eyes of *white* mutant flies is enhanced by mutations in *scarlet*

The *Drosophila* retina is built by about 700 individual units, called ommatidia, which are arranged in a highly stereotypic pattern. Each ommatidium is composed of pigment cells forming a hexagon, which embrace the photoreceptor cells (PRCs). Seven PRCs can be recognized in each section by their rhabdomeres, the photosensitive structure of each cell (Fig. 1E, F). This regular structure is ideally suited to detect even subtle changes, both in tissue structure and in retinal health (for example by reduced number of PRCs).

Flies carrying mutations in the *Drosophila white* (*w*) gene, which lack eye pigments (Fig. 1D), undergo progressive age- and light-dependent retinal degeneration (Ferreiro, Perez et al., 2017, Lee & Montell, 2004), which is enhanced by constant illumination (Bulgakova, Kempkens et al., 2008, Lee & Montell, 2004). To quantify these defects, we defined three phenotypes. (i) Ommatidia with fewer than the normal complement of 7 rhabdomeres (red outlines in Fig. 2A-F). This reflects impaired retinal health, visible at the level of the PRCs. (ii) Density of ommatidia within the tissue. Holes between adjacent ommatidia (highlighted in yellow in Fig. 2A-F) [also called lacunae (Ferreiro et al., 2017)], cause a disruption in the arrangement of adjacent ommatidia. (iii) Intensely-stained aggregates (red arrowheads in Fig. 2B-D), reminiscent of apoptotic bodies. All three phenotypes are indicative of retinal degeneration and a consequence of exposure to high intensity continuous light. For example, holes and lack of rhabdomeres in *w* mutants are not visible when flies are exposed to 12 hours light/12 hours dark (Hebbar et al., 2017). Quantification of rhabdomere number per ommatidium and density of ommatidial units allows us to compare the degree of retinal degeneration in different genotypes following light-exposure. This quantification enabled us to define a spectrum of light-induced histological changes, ranging from mild damage (with qualitative changes as described above), to moderate degeneration (qualitative changes + significant changes to one of the quantified attributes) to severe (qualitative changes + significant changes to both quantified attributes) degeneration.

**Figure 2:**
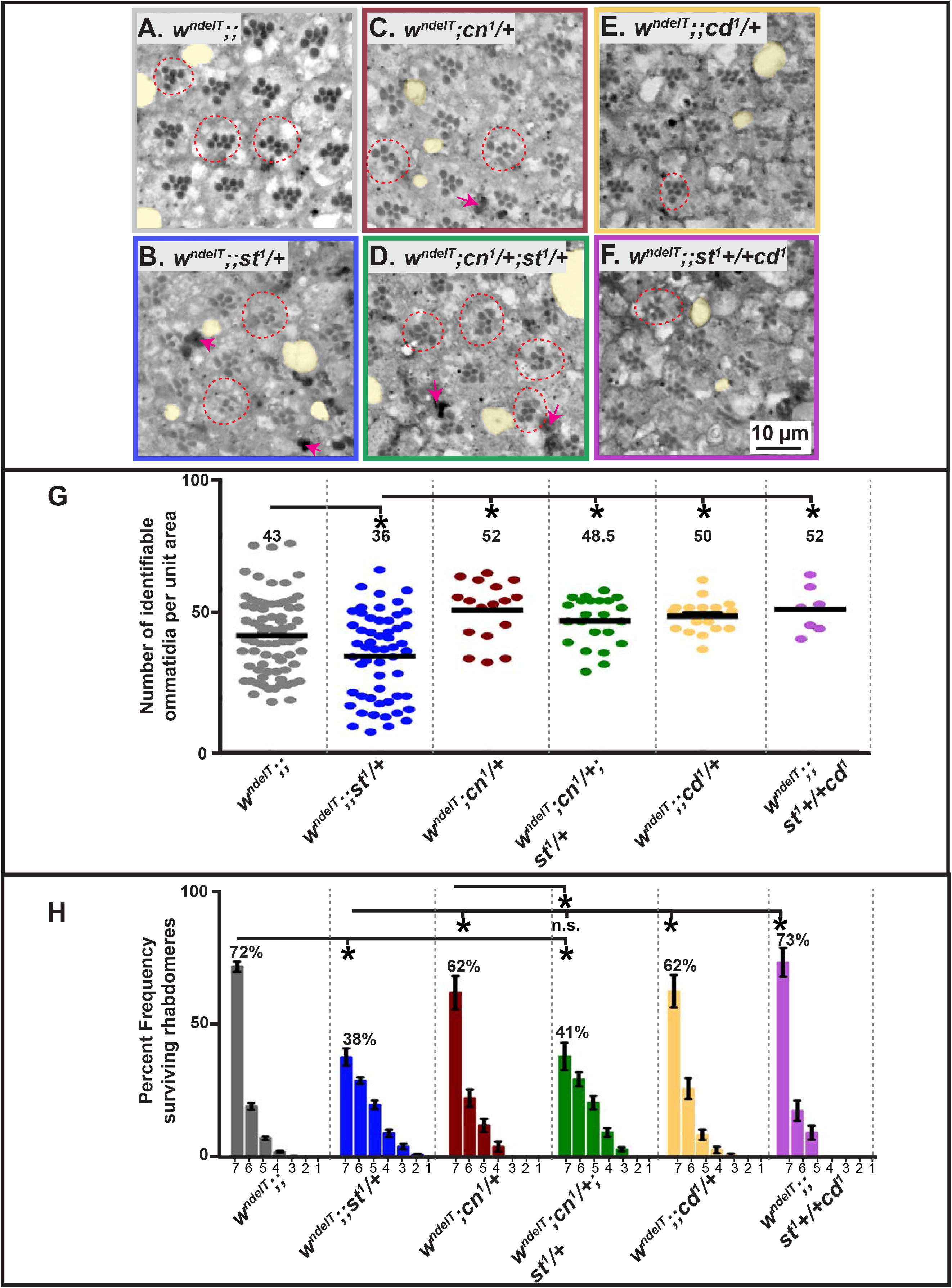
Perturbation of the Kynurenine pathway (KP) modulates retinal damage in *w;;st* mutants. **A-F:** Light microscopic images of toluidine blue-stained, 1μm thin sections of adult fly eyes exposed to continuous, high intensity white light for seven days. Damage attributes are observed such as holes (lacunae) (highlighted in yellow), ommatidia with less than 7 rhabdomeres (red dotted circles), presence of apoptotic debris (intensely stained regions; red arrows). Scale bar as in F. Degenerative attributes appear more pronounced in *w^ndelT^;;st^1^/+* (B; severe) and *w^ndelT^;cn^1^/+;st^1^/+* (D; moderate) than in other genotypes. **G:** Graph comparing the extent of damage following exposure of flies of different genotypes to continuous, high intensity white light for 7 days. Horizontal bars indicate the average number of ommatidia with identifiable rhabdomeres of PRCs, normalized to unit area. Each dot represents an individual count from 1 biological replicate. Numbers above each genotype reflect the average for this population. Only statistically significant differences between genotype pairs, as revealed by ANOVA followed by Bonferroni’s post-hoc test at p<0.05, are indicated by *. **H:** Graph representing mean percent frequency [+ s.e.m.] of ommatidia displaying 1-7 rhabdomeres following continuous, high intensity white light exposure for 7 days. Numbers above the bars indicate the mean value of ommatidia displaying the full complement of 7 rhabdomeres of each genotype. Statistically significant differences between genotype pairs are indicated by * as revealed by ANOVA followed by Bonferroni’s post-hoc test at p<0.05. “n.s” indicates no statistically significant difference.

In all experiments, young flies (2-4 days) were used. We used *w^ndelT^*, a null allele previously designated as *w\** in (Hebbar, Lehmann et al., 2021). In contrast to wild-type eyes, or to eyes mutant for *w^ndelT^* exposed for the same time to less stringent light conditions, e.g. light exposure interspaced with equal intervals of darkness (Hebbar et al., 2021), *w^ndelT^* showed retinal abnormalities upon exposure to continuous, high-intensity light for seven days. First, the retina of light-exposed *w^ndelT^* mutant flies exhibited holes between adjacent ommatidia (Fig. 2A). Nevertheless, the number of identifiable ommatidia per unit area (Fig. 2G) was not significantly different from that in wild-type retinas (Fig. S1A), pointing to largely intact tissue integrity. Second, in *w^ndelT^* mutant retinas, 28% of the ommatidia exhibited less than the full complement of seven rhabdomeres (Fig. 2A, red outlines, and H). Third, there were hardly any large, intensely-stained aggregates in *w^ndelT^* mutant retinas, reminiscent of apoptotic bodies that have been associated with cell death and degeneration in the retina (Johnson, Grawe et al., 2002a). Thus, *w^ndelT^* retinas exposed to high intensity light as applied here exhibited mild signs of light-induced damage.

*Drosophila w* encodes an ATP-binding cassette (ABC) transporter of the ABCG subfamily (Nagao, Juni et al., 2015, Schmitz, Langmann et al., 2001), which is localized in the membrane of the pigment granule-containing compartment, called the lysosome-related organelle (LRO) (Mackenzie, Howells et al., 2000), and is required to transport 3OH-K from the cytoplasm into the LORs (see Fig. 1A). Therefore, we asked whether the mild, light-induced retinal damage in *w^ndelT^* eyes is due to an impaired kynurenine metabolic pathway (KP) rather than to the lack of screening pigments. To address this question, we studied the role of other genes involved in the KP (Fig. 1A) on the retinal phenotype of *w^ndelT^*, since some metabolic intermediates of this pathway, including 3OH-K, have been implicated in other functions, e.g. oxidative stress- and inflammation-induced damage (Colin-Gonzalez et al., 2013).

Flies homozygous mutant for a null allele of *st, st^1^*, exhibit bright-red eyes due to the lack of the brown pigments, the ommochromes (Tearle et al., 1989) (Fig.1C). It should be noted that no retinal degeneration was observed in *st^1^/st^1^* kept under constant light (Suppl. Fig.S1A,B), or under light/dark conditions (Hebbar et al., 2021). Flies homozygous mutant for *w* and *st* (*w;; st/st*) have white eyes similar as *w* due to the absence of all screening pigments. Upon removal of one or both functional copies of *st* in a *w^ndelT^* mutant background, all three aspects of retinal damage of *w^ndelT^* mutants were enhanced (Fig. 2) resulting in severe retinal degeneration. Interestingly, there was no difference in the extent of retinal degeneration with the removal of one or both functional copies of *st* (Fig. S1E-F). First, more lacunae in the retina of *w^ndelT^;;st^1^/+* were observed (compare Fig. 2A with Fig. 2B). This goes along with a significant decrease in the number of identifiable ommatidia per unit area (Fig. 2G) (Hebbar et al., 2021). Second, compared to control, the number of ommatidia with 7 PRCs per section was further decreased, from 72% in *w^ndelT^* to 38% and 32% in *w^ndelT^;;st^1^/+* and *w^ndelT^;;st^1^/st^1^*, respectively (Fig. 2H and Fig. S2F). There was no difference in the degree of light induced damage between male (*w^ndelT^/Y;;st^1^/+*) and female (*w^ndelT^/w^ndelT^;st^1^/+*) flies (data not shown). Third, densely stained pyknotic bodies were abundant, pointing to apoptosis (Fig. 2B, magenta arrows). Exacerbation of light induced damage was also observed when another mutant allele of *w* (*w-RNAi*) was combined with *st^1^* (*w-RNAi;;st^1^*/+) as compared to *w-RNAi* on its own (Fig S1C-D).

Continuous, high intensity light (as used in our experiments), in particular its ultraviolet and blue wavelength component, has been associated with increased oxidative stress and stress-related signaling (Hall, Ma et al., 2018, Lee & Montell, 2004, Roehlecke, Schumann et al., 2013, Roehlecke, Schumann et al., 2011). Therefore, we speculated that it may be oxidative stress and/or stress-related signaling that contribute to the phenotypes described above. In fact, eliminating these shorter wavelengths of light during the course of the constant light exposure abolished damage to retinal tissue in *w^ndelT^* retinas and in *w^ndelT^;;st^1^/+* and *w^ndelT^;;st^1^/st^1^* retinas. None of the genotypes showed signs of tissue damage, as revealed by a comparable number of identifiable ommatidia per unit area in all three genotypes (Fig. S1E). Moreover, almost all (100%, 80%, 99%) ommatidia exhibited 7 rhabdomeres in *w^ndelT^, w^ndelT^;;st^1^/+* and *w^ndelT^;;st^1^/st^1^* eyes, respectively (Fig. S2F). Thus, removal of the stress inducing component of white light is sufficient to not only eliminate the exacerbated degeneration in *w;;st* eyes but also to eliminate the basal level of retinal damage in *w* eyes alone.

Taken together, reduction or loss of *st* function enhanced the severity of the *w^ndelT^* retinal phenotype in the presence of exogenous stress, such as high intensity blue light. This severe retinal degeneration is associated with altered pigment cell homeostasis (preliminary data). In other words, the wild-type function of *st* protects a pigment-less, sensitized *w^ndelT^* retina from degeneration. Since *w^ndelT^, w^ndelT^;;st^1^/+*, *w^ndelT^;;st^1^/st^1^*, *w-RNAi*, and *w-RNAi;;st^1^/+* all have unpigmented (white) eyes, the lack of screening pigments cannot be the cause for the enhancing role of *st*. This raised the question whether the wild-type metabolic roles of *w^+^* and *st^+^* in trafficking tryptophan metabolites in pigment cells (Fig. 1A) (Mackenzie et al., 1999) protect the retina against light-induced damage.

### Mutations in *cinnabar* (*cn*)/KMO modulate the extent of light-induced retinal damage in *w;;st* double mutants

The neuroprotective role of functional *st^+^* in protecting a pigment-less retina from severe light-induced damages is in line with the observation that *st^+^* prevents neurodegeneration in a fly model for Parkinson’s disease (Cunningham et al., 2018). *st*, together with *w*, is involved in the transport of 3OH-K, an intermediate in the kynurenine pathway, from the cytoplasm into LROs (Cunningham et al., 2018, Mackenzie et al., 2000) (Fig. 1A). This led to the suggestion that loss of *st* leads to accumulation of 3OH-K outside the LRO (Cunningham et al., 2018, Mackenzie et al., 2000). In line with this, we hypothesized that excess cytoplasmic 3OH-K levels are responsible for the enhancing effects on the *w^ndelT^* retinal phenotype upon removal of one copy of *st*. To test this hypothesis, we utilized mutations in *cinnabar* (*cn*), the gene that encodes kynurenine monooxygenase (KMO), an enzyme that catalyzes the formation of 3OH-K from Kynurenine (Ghosh & Forrest, 1967, Sullivan, Kitos et al., 1973, Warren, Palmer et al., 1996) (Fig. 1A). We expected that reduced levels of 3OH-K by lowering or removing *cn* activity should attenuate the detrimental effects exerted by loss of *st* on the *w^ndelT^* phenotype. In fact, concomitantly removing one copy of *st* and *cn* in a *w* background (*w^ndelT^;cn^1^/+; st^1^/+*) reduced the severity of the tissue damage observed in *w*; *st^1^/+* retinas. The retinal phenotype of the triple mutant was qualitatively intermediate between the more damaged *w^ndelT^;;st^1^/+* and the less damaged control, *w^ndelT^;cn^1^/+* (compare Fig. 2D with Fig. 2B and C). Ommatidial density was significantly higher in the triple mutant than in *w^ndelT^;;st^1^/+* and very similar to that of *w^ndelT^;cn^1^/+* (Fig. 2G). In contrast, the percentage of ommatidia with seven rhabdomeres in the triple mutant (41%) was similar to that determined for *w^ndelT^;;st^1^/+* (38%) and significantly lower than *w^ndelT^;cn^1^/+* (62%) (Fig. 2H). Therefore, reducing *cn*/KMO attenuates only the ommatidial density phenotype of a *w^ndelT^;;st^1^/+* retina. In contrast, reducing *cn*/KMO does not increase the percentage of surviving rhabdomeres in the triple mutant (Fig. 2H). This supports the notion that modulation of light induced retinal degeneration in these genotypes is reflected primarily in the changes in overall tissue structure (ommatidial density), and the effects on PRCs might be indirect. This phenotype is not allele-specific and could also be observed with *cn^35K^/+*, a hypomorphic allele (Fig. S2A, B). Therefore, reducing *cn*/KMO attenuates the phenotype of a *w^ndelT^;;st^1^/+* retina suggesting a neuroprotective function in line with its suggested role in the biosynthesis of 3OH-K.

### Mutation in *cardinal* (*cd*)/PHS reduces the extent of light-induced retinal damage in *w;;st* double mutants

To further corroborate the hypothesis that 3OH-K abundance plays a role in light-induced retinal degeneration, we studied *cardinal* (*cd*), a gene downstream of *w* and *st* in the KP pathway. *cd* encodes Phenoxazinone synthetase (PHS), an enzyme that participates in the conversion of 3OH-K to the brown pigment in the LRO (Phillips & Forrest, 1970, Phillips et al., 1973) (Fig. 1A). As a result, *cd* mutants have bright red eyes and highly elevated levels of 3OH-K (Phillips et al., 1973, Zhuravlev, Vetrovoy et al., 2020). Therefore, we expected an enhancing effect on the *w^ndelT^* retinal phenotype by removing one copy of *cd*, similar to the effects obtained upon removal of one copy of *st*. However, halving the dose of *cd* in a *white* background (*w^ndelT^;cd^1^/+*) showed less severe light-induced damages than *w^ndelT^;;st^1^/+* retinas (compare Fig. 2E with Fig. 2B). This was reflected by a higher number of identifiable ommatidia per area unit in *w^ndelT^;;cd^1^/+* retinas compared to that of *w^ndelT^;;st^1^/+* (50 vs.36; Fig. 2G) and a higher percentage of ommatidia with the full complement of seven PRCs (62% vs. 38%; Fig. 2H). All features of the mutant phenotype in the triple mutant *w^ndelT^;;st^1^+/+cd^1^* were similar to that of the double mutant *w^ndelT^;;cd^1^/+* and significantly better than the phenotype in *w^ndelT^;;st^1^/+*.

To summarize, removing one copy of *cd*/PHS in a *w* mutant background does not enhance the severity of the *w* mutant phenotype, which is in contrast to the effects observed upon removing one copy of *st* in a *w* mutant background. This result is puzzling, since mutations in both *st* and *cd*/PHS are predicted to have increased 3OH-K levels (Cunningham et al., 2018, Howells & Ryall, 1975, Phillips et al., 1973, Zhuravlev et al., 2020). A notable point is that the Cd protein is localized in the lumen of the LRO (Harris et al., 2011; Fig. 1F) and hence activity of Cd is compartmentalized. Therefore, in *cd* mutants, 3OH-K likely accumulates in the LRO, thus preventing it from exerting its toxicity. In contrast, St is localized in the membrane of the LRO and in conjunction with White, it transports 3OH-K from the cytoplasm to the LRO (Mackenzie et al., 2000).

Overall our genetic experiments of combining *w^ndelT^;;st* with either *cn* or *cd* allowed us to speculate that (1) 3OH-K and/or its metabolites can affect retinal health following light exposure in a sensitized background (such as *w^ndelT^*), and (2) the detrimental consequences of increased abundance of 3OH-K on the *w* retinal phenotype depend on its sub-cellular localization.

### Mutations in *cn/KMO, st*, and *cd/PHS* in a wild-type background have distinct outcomes on the abundances of KP metabolites

To validate the hypothesis that phenotypes described above can be attributed to aberrant levels of 3OH-K and/or its metabolites, we extracted metabolites from the heads of single, double or triple mutants, and identified and quantified them by Mass Spectrometry. We measured Kynurenine (K) and 3-Hydroxykynurenine (3OH-K), the direct metabolites in ommochrome biosynthesis, and Kynurenic Acid (KYNA) and Xanthurenic acid (XA), direct derivatives of K and 3OH-K, respectively (see Fig. 1A). It should be noted that the measurement of metabolites from the different mutants was carried out using extracts of heads of flies kept under normal light/dark conditions to preclude any degeneration. Thus, any change in abundance of these metabolites in a particular genotype could potentially be connected to the susceptibility/vulnerability of that genotype to light stress.

The data obtained from the measurements of the metabolites in the different eye color mutants, performed in a wild-type background, are listed in Table 1 and schematized in Suppl. Fig. S3. Major results can be summarized as follows.

**Table 1:**
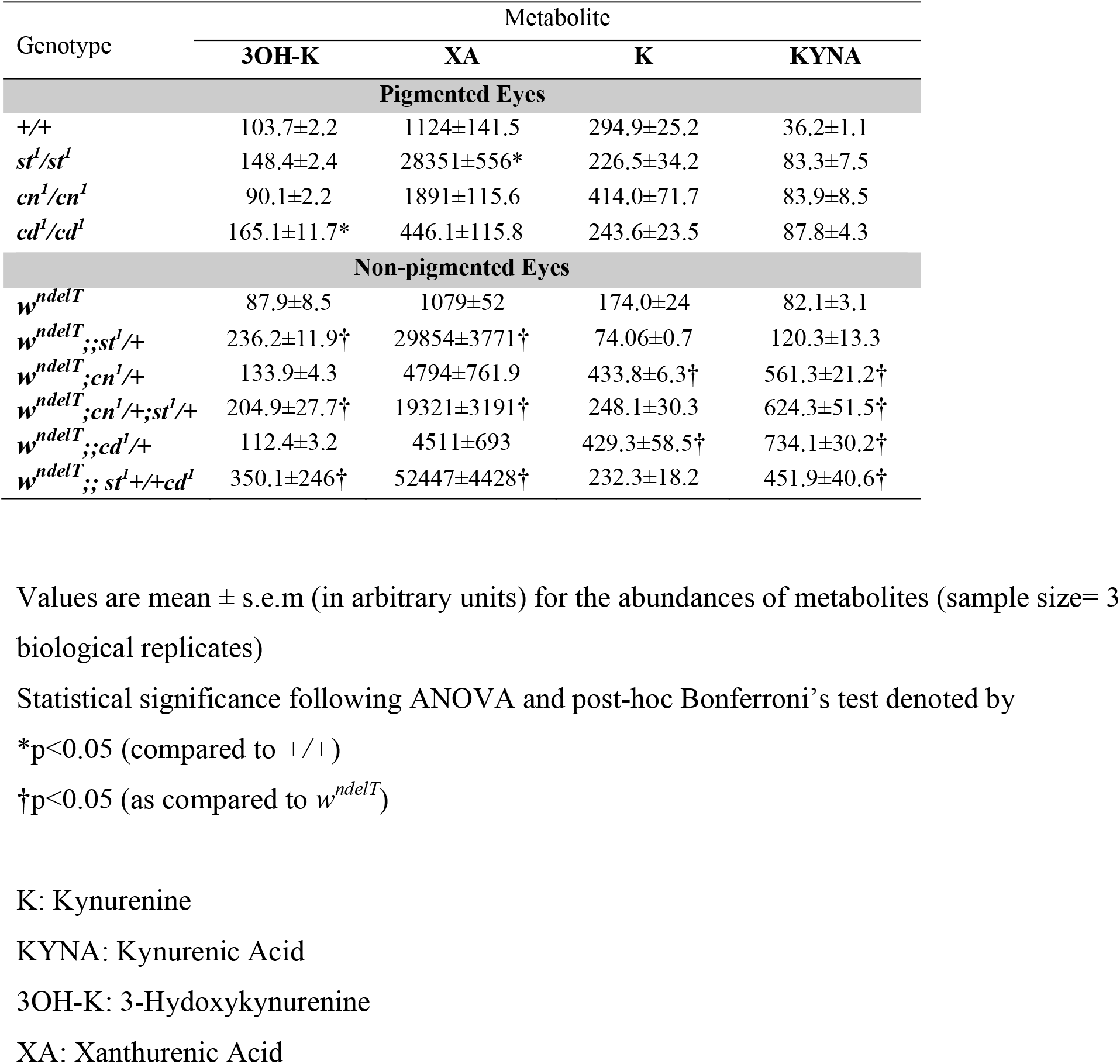
Metabolite abundance across genotypes

First, removing both copies of either *cn* or *st* in an otherwise wild-type background (red eyes) exhibited no change in the levels of 3OH-K of adult heads (Suppl Fig. S3A’, lanes 2 and 3, respectively).

Second *cd* mutants showed significantly higher 3OH-K levels than the wild-type control, as expected (Suppl Fig. S2 A’, lane 4).

Third, 3OH-K was significantly increased in *st^1^/st^1^* and *cd^1^/cd^1^* retinas compared to *cn^1^/cn^1^* (Suppl Fig. S2A’, lanes 2, 4 versus 3 respectively).

Fourth, XA, a metabolite of 3OH-K (see Fig. 1A) was significantly higher in *st^1^/st^1^* mutant retinas, but not in *cd^1^/cd^1^* retinas, when compared to wild-type (Suppl Fig. S2B’, lanes 2, 4 versus lane 1).

Fifth, no significant changes were observed for K and KYNA in the genotypes examined when compared to wild-type (Table 1 and Suppl Fig. S3 C-D’).

Fig. 3A’ illustrates the results by showing the ratio of precursor/product, i.e. [K]/[3OH-K]. In agreement with data from another allele, *cn^3^* (Campesan et al., 2011, Maddison, Alfonso-Nunez et al., 2020, Sasaki, Nishimura et al., 2021), and as expected from the lack of an enzyme (KMO) that converts K to 3OH-K, this ratio is significantly increased in *cn^1^* mutant eyes. On the other hand, no change in ([K]/[3OH-K]) was observed in *st* mutants. This is consistent with previous reports, which explained the lack of increased 3OH-K in heads of *st* mutant adults by its excretion via Malpighian tubules at the end of pupal development (Howells & Ryall, 1975, Phillips et al., 1973). Interestingly this elimination of 3OH-K is not observed in *cd*/PHS mutants [also in (Howells, Summers et al., 1977)], suggesting that here compartmentalization and/or another alternate mechanism sequesters 3OH-K.

**Figure 3:**
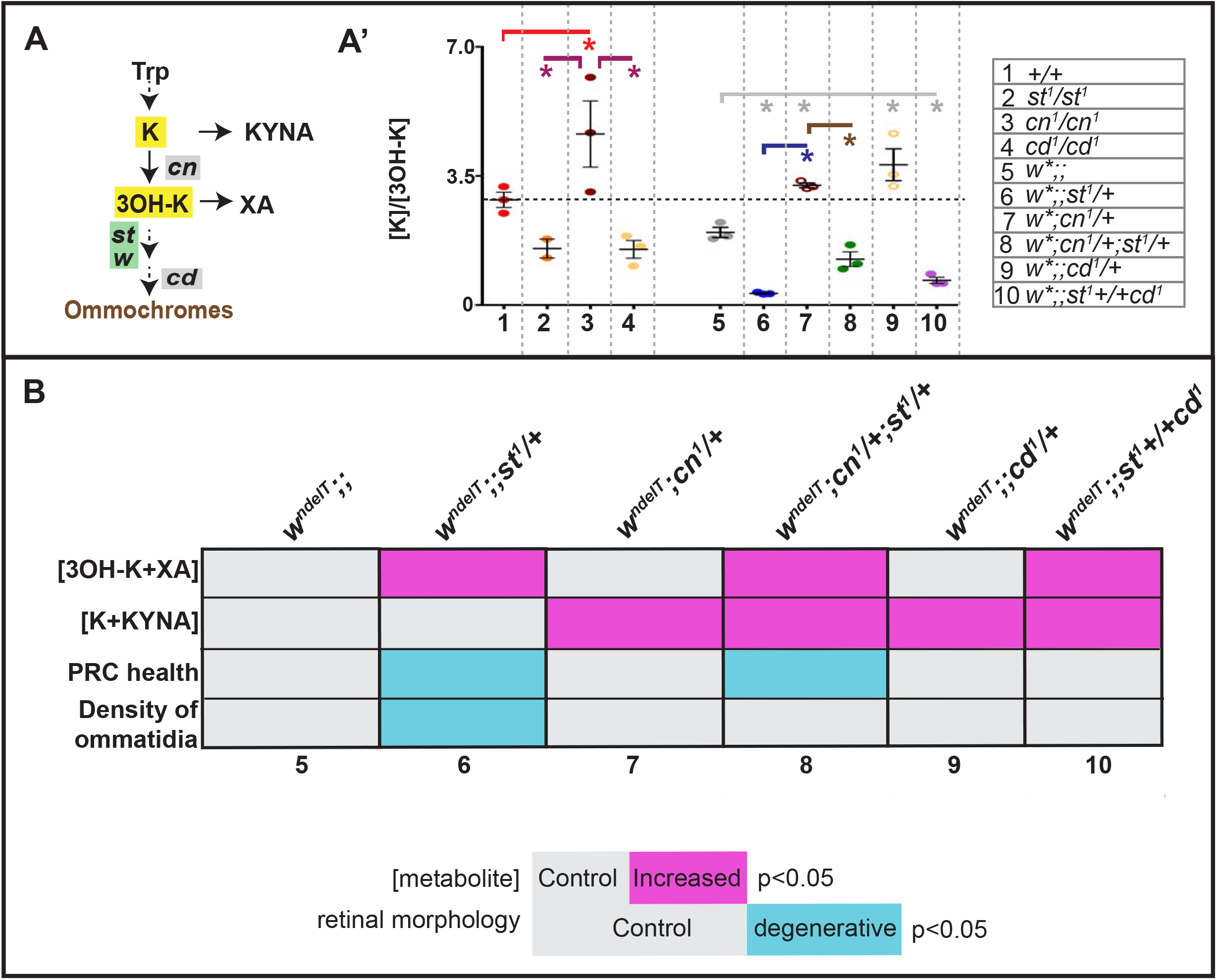
Increased 3OH-K abundance is associated with extensive light induced retinal damage. **A:** Schematic outline of relevant steps in the Kynurenine pathway (KP) and brown pigment biosynthesis. Genes used in this study are indicated. **A’:** Graph depicting the ratio of the abundances of Kynurenine and 3-hydroxykynurenine [K]/[3OH-K] (highlighted in yellow in the corresponding pathway shown in A-D) from M.S. measurements of head extracts of flies kept under physiological/non degenerative light conditions. Numbers on X-axis correspond to the genotypes listed in the table on the right. Each colored dot represents an individual biological replicate of 10 pooled heads. Statistical comparisons between genotype pairs are indicated with solid lines (red line compared to +/+, magenta line compared to *cn^1^/cn^1^*, grey line compared to *w^ndelT^*, blue line compared to *w^ndelT^;;st^1^/+*, brown line compared to *w^ndelT^;cn^1^/+*) with * revealed by ANOVA followed by Bonferroni’s post-hoc test at p<0.05. **B:** Tabular summary of various parameters (horizontal lines) with different genotypes (columns 5-10, corresponding to genotypes shown in the table in A’). The parameters include metabolite levels (top two) and retinal health indicators (bottom two). For metabolite levels, grey colored cells imply no difference from controls (*w^ndelT^*), cells colored magenta imply significantly higher metabolite levels. For retina health indicators, grey colored cells imply no difference from controls (*w^ndelT^*), cells colored cyan imply significantly reduced ommatidial density (number of identifiable ommatidia per unit area) or PRC health (percent frequency of surviving rhabdomeres). Statistical significances were calculated with ANOVA followed by Bonferroni’s post-hoc test at p<0.05.

In conclusion, our measurements of KP metabolites in pigmented eyes provide a biochemical characterization of the KP as shown in Fig. 1A. Further, these data support the idea that mutations in *st* and *cd*/PHS have different outcomes on the KP.

### Increased abundance of 3OH-K in *w;;st* mutants is associated with increased susceptibility to retinal damage

Next, we determined the effects of removing single copies of genes of the KP (*cn*/KMO, *st*, *cd*/PHS) on the different KP metabolites in a *w* mutant background, to correlate with changes in mutant phenotype.

First, *w^ndelT^* on its own displayed no significant changes in any of the metabolites when compared to +/+ (Table 1; Suppl Fig. S3).

Second, significantly higher 3OH-K and XA levels compared to *w^ndelT^* were observed in the double mutant *w^ndelT^;;st^1^/+* and in the triple mutants *w^ndelT^;cn^1^/+; st^1^/+* and *w^ndelT^;;st^1^ +/+ cd^1^* (Table 1; Suppl. Fig. S3A’, B’, lane 6 versus lane 5, lane 8 versus lane 5, and lane 10 versus lane 5, respectively).

Third, K and KYNA abundances were unaltered in *w^ndelT^;;st^1^/+* mutant as compared to *w^ndelT^* (Suppl. Fig. S3C’ and D’, lane 6 versus lane 5).

Fourth, significantly higher levels of K and KYNA were detected in *w^ndelT^;cn^1^/+* and *w^ndelT^;cn^1^/+;st^1^/+* compared to *w^ndelT^* and *w^ndelT^;;st^1^/+*, respectively (Suppl. Fig. S3C’ and D’, lane 7 versus lane 5, and lane 8 versus lanes 6, respectively).

Fifth, a significant increase in KYNA levels was observed in *w^ndelT^;;cd^1^/+* and *w^ndelT^;;st^1^+/+cd^1^* as compared to *w^ndelT^* (Table 1 and Suppl. Fig S3 D’, lane 9 versus lane 5 and lane 10 versus lane 5).

Overall, these measurements corroborate the idea that the KP is sensitized by mutations in the *w* gene. This is further demonstrated by examining the ratio of K/3OH-K (Fig. 3A). Reducing one component of the KP (either *cn*/KMO, or *st*) in a *w* genetic background results in more pronounced changes in the ratio of K/3OH-K as opposed to its reduction in a wild-type background alone (Fig. 3A: in the case of *cn*, lane 1 versus lane 3 and lane 5 versus 7 and in the case of *st*, lane 1versus lane 2 and lane 5 versus lane 6,).

Do these metabolite abundances (measured in physiological conditions) correlate with the susceptibility of the retina to light exposure? To better visualize the associations of altered metabolite abundances with retinal health, we designed a color-coded table (Fig. 3B), which revealed important trends.

First, increased 3OH-K+XA abundance is linked with severe and moderate retinal phenotypes, as in *w^ndelT^;;st^1^/+* and *w^ndelT^;cn^1^/+;st^1^/+*, respectively (Fig. 3B, columns 6 and 8). Accordingly, a healthy retina correlates with no changes in 3OH+XA abundance (Fig. 3B, columns 5, 7, 9).

Second, the moderate retinal phenotype in *w^ndelT^;cn^1^/+;st^1^/+* despite increased 3OH-K+XA can be explained by concomitant increased K+KYNA abundance (Fig. 3B, column 8), which most likely counteracts the detrimental effects of increased levels of 3OH-K and XA. However, differences in the extent of the rescue between *w^ndelT^;cn^1^/+;st^1^/+* and *w^ndelT^;;st^1^ +/+ cd^1^* cannot be explained by these metabolite abundances alone (Fig. 3B, column 8 versus 10). The moderate degeneration in case of *w^ndelT^;cn^1^/+;st^1^/+* might stem from a separate role of *cn*/KMO in mitochondrial regulation (Maddison et al., 2020) independent of metabolite levels.

In conclusion, these results connect increased levels of KYNA and K with better retinal health, most likely by counteracting the detrimental effects of increased levels of 3OH-K and XA.

### Feeding 3OH-K is sufficient to enhance light-induced retinal damage in a *w^ndelT^* sensitized genetic background

Our data suggest that 3OH-K metabolites are “toxic” and sensitize the *w^ndelT^* retinal tissue to light-induced damage (as in *w^ndelT^;;st^1^/+*), while KYNA exerts a “protective” function (as in *w^ndelT^;cn^1^/+;st^1^/+*). To further corroborate this assumption, we fed animals with “protective” KYNA (1mg/ml) or “toxic” 3OH-K (1mg/ml) and evaluated retinal health upon light exposure. Similar amount of these metabolites has been previously used to demonstrate their importance in neurodegenerative mutants (Campesan et al., 2011). The feeding regimen used here was sufficiently effective for the incorporation of the metabolites into eyes. For example, feeding 3OH-K (1mg/ml) to *cn^1^/cn^1^* mutants, which cannot synthesize 3OH-K and hence develop bright-red eyes due to an impaired ommochrome pathway, was sufficient to restore the complete brown pigment biogenesis pathway and change the eye color to wild-type (Suppl Fig. S4A). This treatment had no consequences on retinal morphology following light exposure (Fig. 4A, B), consistent with our earlier observations that KP disruption does not impair retinal morphology upon light exposure in the wild-type background/pigmented eyes.

**Figure 4:**
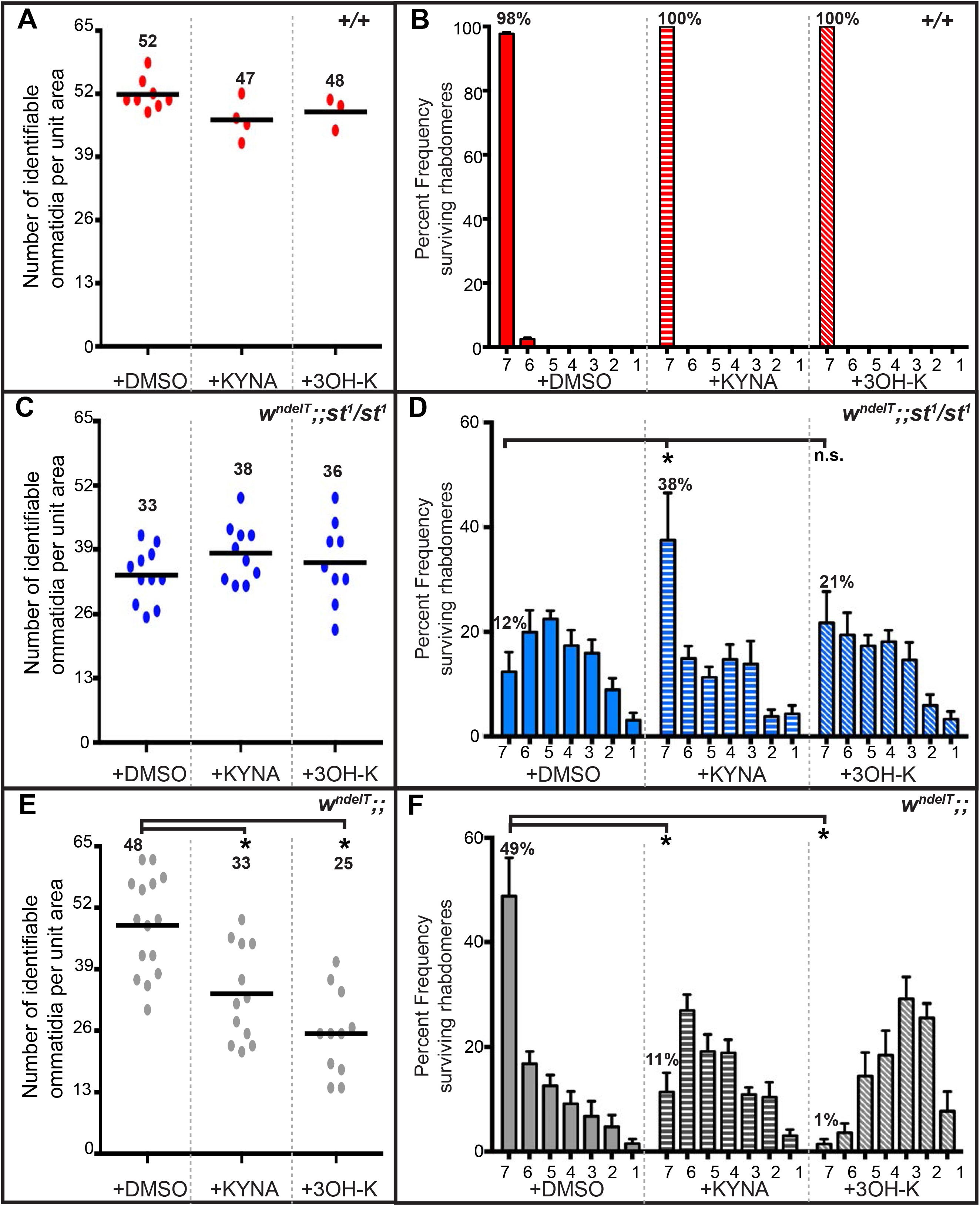
Effect of light induced damage upon feeding 3OH-K or KYNA to flies of different genotypes in sensitized genetic backgrounds. **A, C, E.** Graphs comparing the extent of retinal damage following exposure to continuous white light in +/+ (red; A), *w^ndelT^;;st^1^/st^1^* (blue; C) and *w^ndelT^* (grey; E) flies raised on control food (+DMSO) or food supplemented with either 1mg/ml KYNA or 1mg/ml 3OH-K. Horizontal bars indicate the average number of identifiable ommatidia normalized to unit area. Each dot represents an individual count from 1 biological replicate. Number indicates the mean value for each condition. Statistically significant differences between genotype pairs are indicated by * as revealed by ANOVA followed by Bonferroni’s post-hoc test at p<0.05. **B, D, F.** Mean percent frequency of ommatidia displaying 1-7 rhabdomeres [± s.e.m.] upon exposure of flies to continuous, high intensity white light for 7 days. +/+ (red; B), *w^ndelT^;;st^1^/st^1^* (blue; D) and *w^ndelT^* (grey; F) flies were raised on control food (+DMSO) or on food supplemented with either 1mg/ml KYNA (bars with horizontal lines) or 1mg/ml 3OH-K (bars with slanted lines). Numbers above the bars indicates the mean value of ommatidia displaying the full complement of 7 rhabdomeres. * indicates statistical significance for the pair as revealed by ANOVA followed by Bonferroni’s post-hoc test at p<0.05. “n.s” indicates no statistically significant difference.

In contrast, feeding “protective” KYNA to *w^ndelT^;;st^1^/st^1^* animals reversed PRC damage observed in control animals (fed with DMSO). 38% of the ommatidia of flies fed with KYNA showed seven rhabdomeres compared to 12% observed in the feeding control (Fig. 4D). Thus, feeding KYNA phenocopied the triple mutant *w^ndelT^;cn^1^/+; st/+^1^*. However, this treatment did not significantly increase the number of identifiable ommatidia per unit area (a measure for tissue integrity) (Fig. 4C). Feeding “toxic” 3OH-K to *w^ndelT^;;st^1^/st^1^* flies did not further exacerbate light-induced retinal damage in terms of ommatidial integrity (Fig. 4C) and PRC damage (Fig. 4D). In contrast, feeding 3OH-K in another sensitized background (*w^ndelT^*) resulted in significantly more damage compared to the feeding control (with DMSO). The number of identifiable ommatidia per unit area decreased from 48 to 25 (Fig. 4E) and the percentage of ommatidia containing seven rhabdomeres was significantly reduced upon feeding 3OH-K, from 49% in the control to 1% (Fig. 4F). Finally, in contrast to our expectation, feeding *w^ndelT^* animals with KYNA increased the severity of both read-outs of retinal integrity (Fig. 4E, F).

Taken together, dietary increase of KYNA protected the *w^ndelT^;;st^1^/st^1^* retina from light-induced PRC damage, and hence provides a molecular explanation for the phenotype of the triple mutant. 3OH-K feeding, on the other hand, enhanced the phenotype of *w^ndelT^* flies, thus phenocopying *w^ndelT^;;st^1^/+* mutant retinas (Fig. 2), which have increased 3OH-K levels (Table 1). Overall, these data strengthen our earlier conclusions that increased 3OH-K (or increased 3OH-K +XA) makes a retina more susceptible to light-damage. On the other hand, increased KYNA (or increased K + KYNA) is protective in a sensitized background.

### Free 3OH-K, but not protein-bound 3OH-K, is connected to susceptibility to light-induced degeneration

Although both 3OH-K and K+KYNA levels are increased in in *w^ndelT^;cn^1^/+;st^1^/+* and in *w^ndelT^;;st^1^ +/+ cd^1^*, only the former shows a mutant retinal phenotype (compare column 4 and 6 in Fig. 3). This raised the question of the status of free and protein-bound 3OH-K (PB-3OH-K) in the context of the genotypes studied here, since 3OH-K can exist in a protein-bound form in flies (Zhuravlev et al., 2020) and in humans (Korlimbinis & Truscott, 2006). To test this, we determined the levels of PB-3OH-K in head lysates of flies raised in non-degenerative light conditions, both in wild-type (+/+) and in *w^ndelT^* mutant background. PB-3OH-K was detected in head lysates of +/+, and, in lower amounts, in head lysates of *cd^1^/cd^1^* flies (Fig. 5A, lane 1 versus lane 3) as reported previously (Zhuravlev et al., 2020). Increased PB-3OH-K was also detected in +/+ flies fed with 3OH-K (Suppl Fig. S4B). In stark contrast, PB-3OH-K was not detected in head lysates of *st^1^/st^1^* (Fig. 5A, lane 2), nor in all genotypes with *w* in the background, namely *w^ndelT^, w^ndelT^;;st^1^* /+, and *w^ndelT^;;cd^1^/+* (Fig. 5A, lanes 4-6 respectively) and *wRNAi* (Fig. 5B, lane 7). These results suggest that 3OH-K is bound to proteins only upon its transport into LROs (Fig. 5C) by functional White/Scarlet (Mackenzie et al., 2000). PB3OH-K was also decreased upon the concomitant knockdown of *st* and *w* by RNAi (Fig. 5B lane 9) as opposed to knockdown of *w* alone (Fig. 5B, lane 8).

**Figure 5:**
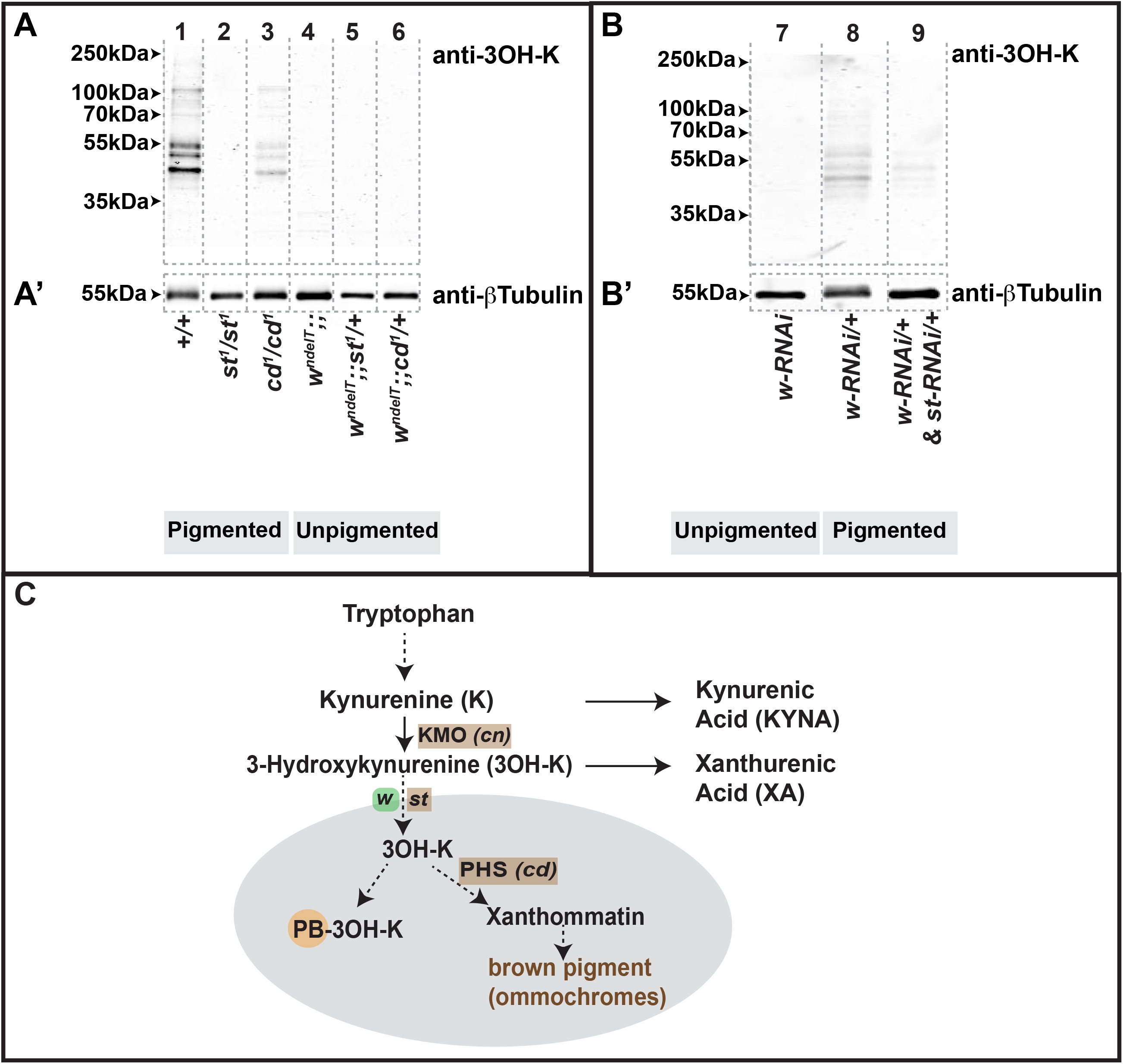
Perturbation of the Kynurenine pathway affects the amount of protein-bound 3OH-K. **A-B’:** Western blots of protein extracts from heads of flies of different genotypes raised on normal food under physiological/non-degenerative conditions. Blots were tested with antibodies against 3OH-K (A, B) and β-Tubulin as loading control, run on parallel gels (A’, B’). Numbers on the top of the lanes correspond to the genotypes listed below. Positions of standard molecular weight markers are indicated by arrowheads. Vertical, grey dotted lines outline the lanes. **C:** A cartoon depicting selected steps of the Kynurenine pathway. Kynurenine 3-monooxygenase (KMO) and Phenoxazinone synthetase (PHS) are enzymes encoded by *cinnabar* (*cn*) and *cardinal* (*cd*), respectively. White and Scarlet, encoded by *white* (*w*) and *scarlet* (*st*), respectively, compartmentalize free 3OH-K through their transporter activity into LROs (grey). The schematic represents the emerging hypothesis from the experiments shown in A-B’ that 3OH-K can be bound to proteins (PB-3OH-K) only following its transport into LROs by White and Scarlet

On the other hand, in *cd* mutants, free 3OH-K is still transported into LROs, but cannot been utilized for the formation of brown pigment (Fig. 5C). However, it is sequestered by conjugation with proteins as evidenced by PB-3OH-K in head lysates of *cd^1^/cd^1^* (Fig. 5 A’, lane 3). Further, it should be noted that PB-3OH-K was only detected in wild-type and *cd^1^/cd^1^*, genotypes, both of which do not exhibit retinal damage when exposed to light (Suppl Fig. S1A-A’). But why do *st* mutant eyes, with no visible PB-3OH-K levels, not exhibit light induced damage? Residual red pigmentation in *st* mutants absorbs light and shields the retina from high intensity light. When this residual pigmentation is removed as in *w* mutants, which also by themselves do not show increased PB-3OH-K, high intensity light induces retinal damage. Thus, the compartmentalization and conjugation of 3OH-K to proteins suggests a mechanism allowing to sequester and subsequently “detoxify” the effects of free 3OH-K.

Overall, data shown here allow the conclusion that increase of free 3OH-K is detrimental for retinal health in *Drosophila*, in particular in a sensitized background (*w*). More importantly, they unravel the importance of compartmentalization of the KP metabolites as a protective mechanism achieved by sequestering of 3OH-K, either via the formation of xanthommatin in LROs by *cd*/PHS, or via the conjugation of free 3OH-K to proteins.

## Discussion

Results presented here underscore the importance of metabolites of the Kynurenine pathway (KP), one of the catabolic pathways for tryptophan, in neurodegeneration. Through the use of a series of fly mutants that perturb distinct steps of the KP (Fig. 5C), metabolite analysis, and dietary supplementation, we propose two major mechanisms by which the net outcome of the KP modulates retinal degeneration. First, although increased 3OH-K levels can be detrimental, more important is the balance between toxic 3OH-K and protective KYNA that impacts on the degree of degeneration. In other words, modulating the balance of KP metabolites in favour of increased KYNA protects the retina. Second, 3OH-K only worsens light-induced retinal degeneration when present as free 3OH-K, but not in its protein-bound form (PB-3OH-K). This study now brings to the forefront this previously understated aspect of the KP in the context of tissue health. Sequestering of 3OH-K, in protein-bound form or by compartmentalisation, represents a mechanism to mitigate its toxicity in neurodegeneration. This becomes relevant when considering the KP as a therapeutic target in neurological disorders (Dantzer, 2017, Stone & Darlington, 2013).

The toxicity of 3OH-K stems from its ability to act as a pro-oxidant that auto-oxidizes (non-enzymatic) itself, followed by oxidation of other proteins or cellular DNA [reviewed (Thomas & Stocker, 1999)]. On the other hand, 3OH-K was shown in different contexts to act as a potent scavenger of radicals, such as reactive oxygen species (ROS). The non-enzymatic oxidation of 3OH-K leads to formation of hydroxanthommatin radicals, which are then dimerized to form xanthommatin (Phillips & Forrest, 1970, Wiley & Forrest, 1981). Hence, 3OH-K has the potential to act both as an oxidant, producing more ROS, and as antioxidant, depending on its concentration and the overall oxidizing environment in cells (Zhuravlev, Vetrovoy et al., 2018). The link of 3OH-K and ROS is particularly important in neuronal cells, including those in the brain and the retina, which are metabolically very active and therefore more vulnerable to hyperoxidation. Brains of *st* mutants have elevated ROS levels, which could be reduced upon lowering 3OH-K levels (Cunningham et al., 2008), suggesting that 3OH-K exacerbates ROS production. Preliminary data allowed us to speculate that a similar increase of ROS in the sensitized *w; st/+* retina makes this tissue more prone to degeneration under light stress. This assumption is supported by the observation that degeneration only takes place upon exposure to high intensity light, in particular light containing UV- and blue-wavelengths (this study). These shorter wavelengths enhance ROS production (Chen, Hall et al., 2017) and result in an increased oxidative stress response (Hall et al., 2018). Hence, flies with underlying genetic perturbation of the KP leading to increased 3OH-K are more vulnerable to retinal degeneration when exposed to increased ROS-producing conditions such as high intensity continuous light.

Apart from ROS production, the KP affects many other cellular processes. For example, KP metabolites are regulators of zinc storage in *Drosophila* Malpighian tubules (Garay, Schuth et al., 2022). KP genes such as *cn*/KMO also play a role in mitochondrial dynamics, independent of metabolite abundances (Maddison et al., 2020). Likewise, retinal health and homeostasis depend on several factors/processes, dysregulation of which, including defects in rhodopsin (Xiong & Bellen, 2013), lipid homeostasis (Liu, Zhang et al., 2015, Muliyil, Levet et al., 2020, Raghu, Yadav et al., 2012, Van Den Brink, Cubizolle et al., 2018) and iron storage (Chen, Lin et al., 2016) to name a few, have been implicated in retinal degeneration. Future analyses are necessary to unravel interactions between these cellular processes and the KP in retinal homoestasis.

Results presented here provoke two important questions:

First, *why does a white (w) genetic background influence KP metabolite abundances differently in cn, st, or cd mutants?* This is puzzling because loss of a functional copy of *w*, on its own, does not exhibit any significant changes in KP metabolites. Yet, *w* in combination with *cn, st or cd mutants*, results in pronounced changes to KP metabolite abundances, more than when *cn, st or cd mutants* are in a wild-type background. Unexpected changes in KP metabolites due to a *w* genetic background are, however, not unique to our study, and were also observed upon modifying the dosage of *w* (Campesan et al., 2011). Since measurements as presented here only depict a snapshot of the abundances of these KP metabolites, we cannot exclude an indirect influence as a result of loss of *whit*e on the turnover of KP metabolites in these genotypes. These indirect effects could, for example, stem from the role of *white* in the import of the precursor Tryptophan (Sullivan, Bell et al., 1980), or in its role in the metabolism of tetrahydrofolate (Sasaki et al., 2021) or sphingolipid degradation (Wang, Lin et al., 2022).

Second, *why does increased 3OH-K not always aggravate the retinal phenotype in a sensitized w background?* Fly models for Parkinson’s disease (Cunningham et al., 2018), Huntington’s disease [HD; (Campesan et al., 2011)] and retinal degeneration (this work) reinforce a previously reported connection between neurodegeneration and an imbalance in toxic and protective KP metabolites (Lovelace, Varney et al., 2017, Maddison & Giorgini, 2015, Zinger, Barcia et al., 2011). These models also allowed to establish a causal relationship between an imbalance of metabolites of the KP and the severity of a mutant phenotype. Therefore, the consequence of a disrupted KP in the context of neuronal degeneration should not only take into account changes in the absolute amounts of individual metabolites, but also any imbalances between toxic (3OH-K and XA) and protective (KYNA and K) metabolites as well as their possible conversion to non-toxic moieties (PB-3OH-K). For example, removing one copy of *st* in a *w* mutant background results in elevated 3OH-K levels and severe retinal degeneration. Removing one copy of *cd*/PHS in this background does not affect retinal health despite similarly increased 3OH-K levels, due to concomitant increased K+KYNA abundance. This suggests that increased KYNA levels (reported here) counterbalance the increase in 3OH-K. Elevated KYNA can be protective since it has the potential to neutralize ROS and reduce the oxidative stress burden (Lugo-Huitron, Blanco-Ayala et al., 2011). Indeed, we could show that feeding KYNA in a sensitized (*w;;st*) background reduces the severity of degeneration. Future studies on the turnover of KP metabolites should be assayed to better understand the upregulation of KYNA in mutant combinations of *w* with *cn*/KMO or *cd*/PHS.

Another mechanism that might explain the lack of toxicity of increased 3OH-K, which have not been considered previously in the fly models, is sequestering of 3OH-K by its conjugation to proteins. This process depends on the transport of 3OH-K into LROs by the W/St transporters (Fig. 7B). *cd*/PHS mutants (in a wild-type background) display increased 3OH-K levels. Although *cd* mutants are defective in conversion of free 3OH-K to xanthommatin, they nevertheless form PB-3OH-K, though in lower amounts, thus reducing the detrimental effects of free 3OH-K. This result could also explain a recently published observation that *cd*/PHS mutants show age-related deficits in courtship songs and memory (Zhuravlev et al., 2020). Furthermore, it might help clarify the contradictory data connecting increased abundance of 3OH-K to neurodegeneration in some studies, and to survival in others.

While conjugation of 3OH-K to proteins reduces free, toxic 3OH-K, at the same time this modification can alter the function of proteins and can convert them to harmful variants. For example, 3OH-K conjugation to Aβ peptides and peptides derived from α-synuclein was suggested to be related to neurodegeneration (Capucciati, Galliano et al., 2019). In contrast, 3OH-K conjugated to the human eye lens protein (Staniszewska & Nagaraj, 2005) and was detected in normal aged human lenses but not in damaged (cataract) lenses (Korlimbinis & Truscott, 2006). In this case, 3OH-K-based protein modifications may actually protect the retina from UV-light induced damage, and is probably not the cause of age-related eye damage (Korlimbinis & Truscott, 2006). Data obtained here in the fly retina also support the notion that PB-3OH-K is not harmful, but rather quenches free toxic 3OH-K. Formation of PB-3OH-K is common to conditions that do not display any light induced degeneration, including wild-type, *cd*/PHS mutants, and wild-type flies fed with 3OH-K.

In contrast, is the absence of PB-3OH-K associated with increased susceptibility to retinal damage under conditions of light stress? This is indeed the case for mutations in the ABCG transporter/*w*. Mutations in *w*, or *w* in any genetic combination studied here, displayed almost no PB-3OH-K and were indeed susceptible to retinal damage of varying severities following light exposure. This suggests that free 3OH-K can be toxic when not converted to xanthommatin or conjugated to proteins. This non-sequestered/free 3OH-K, albeit in the physiological abundance range, might underlie the mild phenotypes in *w* retinas when exposed to light, and hence constitutes a sensitized genetic background. In this context, it should be noted that PB-3OH-K is also absent in *st* retinas, resulting in non-sequestered 3OH-K, yet *st* retinas in a wild-type background do not degenerate. When exposed to high intensity light, the *st* retina is presumably shielded from intense light exposure due to the presence of the red shielding piments. The complete absence of red and brown pigments in *w* therefore make it a more susceptible background when compared to *st* in our study.

In the future, conjugation of 3OH-K to proteins may be considered as a potential therapeutic strategy to be tested in case of other neurodegenerative diseases with KP disruption. It would also be relevant to understand the process and sub-cellular location in which the covalent addition of 3OH-K to proteins occurs. PB-3OH-K was detected by our method in head extracts of *cd*/PHS mutants but not in *w* or *st* mutants. This revealed that 3OH-K conjugation to proteins occurs only once it is compartmentalized into LROs (Fig. 5C). In vitro experiments have indicated that 3OH-K is most likely attached to cysteine residues of proteins (Korlimbinis & Truscott, 2006). However, the identity of the target proteins and the functional consequence of their modification still remain elusive.

Put together, this work highlights a novel function for genes involved in the KP in regulating *Drosophila* retinal health, which is uncoupled from their role in ommochrome formation. The results underscore the importance of compartmentalization of a KP metabolite, 3OH-K, and its quenching to non-toxic PB-3OH-K, in regulating tissue homeostasis.

---

**Supplemental Figure S1:**
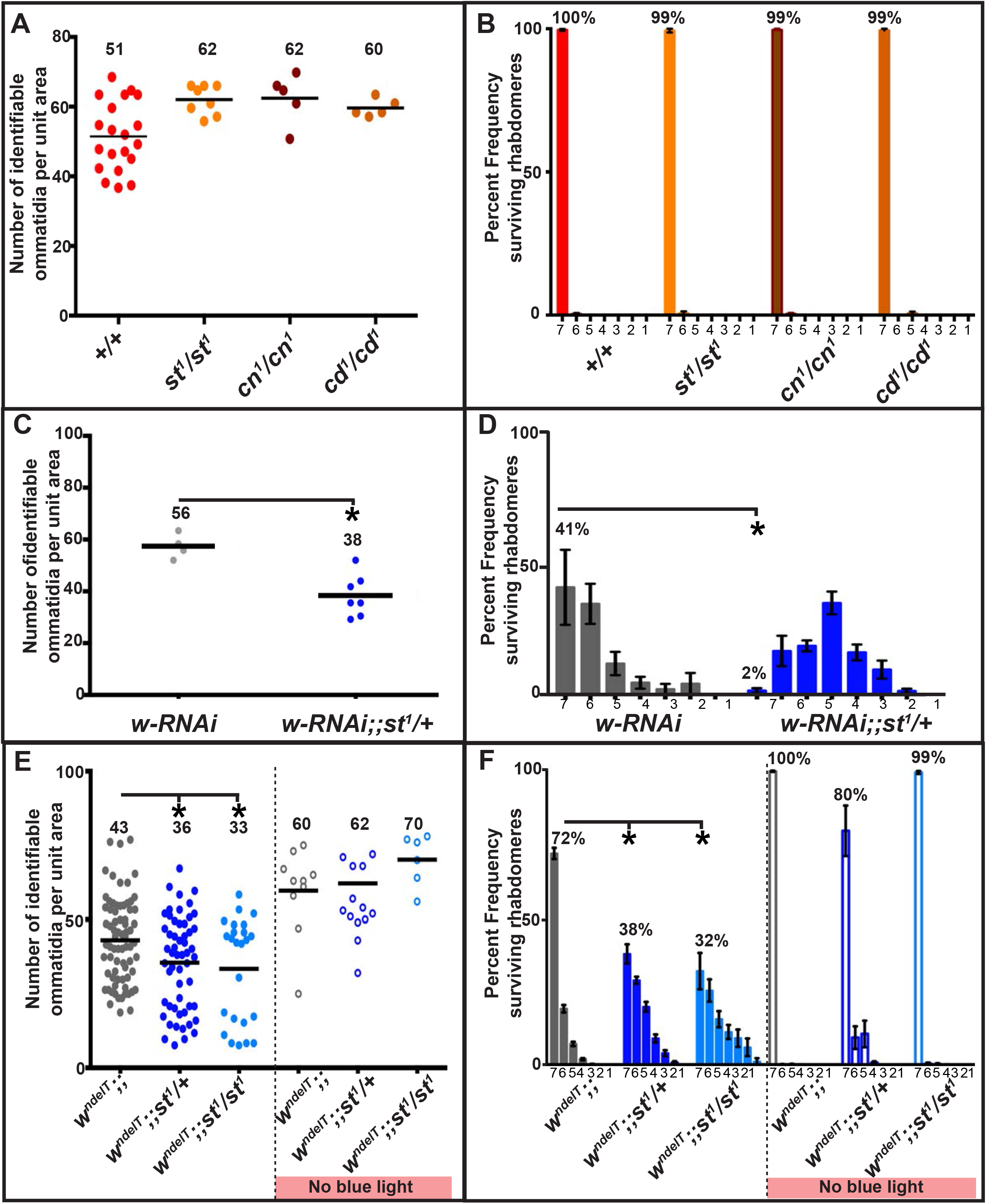
The effect of pigmentation and shorter wavelength light on the extent of light-induced retinal damage in different genotypes. In A, C, E: Horizontal bars indicate the average number of ommatidia with identifiable rhabdomeres normalized to unit area. Each circle represents an individual count from 1 biological replicate. Number indicates the mean value for each condition. Only statistically significant differences, between genotype pairs as revealed by ANOVA followed by Bonferroni’s post-hoc test at p<0.05, are indicated by * (or Unpaired t-test p<0.05, in C). In B, D, F: Percent frequency of ommatidia [± s.e.m.] displaying 1-7 rhabdomeres upon 7 days of continuous, high intensity, light exposure. Numbers above the bars on top of each bar indicates the mean value of ommatidia displaying the full complement of 7 rhabdomeres. No statistically significant differences were observed between genotype pairs as revealed by ANOVA followed by Bonferroni’s post-hoc test at p<0.05 (or Unpaired t-test p<0.05, in D). In A-F, flies were exposed to high intensity, continuous light for 7 days. No damage is observed in a *w^+^*, pigmented background (A.B). Knocking-down *w* (*w-RNAi*) and removing of one copy of *st (w-RNAi;;st^1^/+*) results in retinal damage as compared to *w-RNAi* alone (C, D). Exposure to continuous, high intensity light lacking the short, blue wavelength component for 7 days (unfilled circles in E, unfilled bars in F) does not induce retinal damage of the indicated genotypes as compared to light with the short, blue wavelength component (filled circles in E and filled bars in F).

**Supplemental Figure S2:**
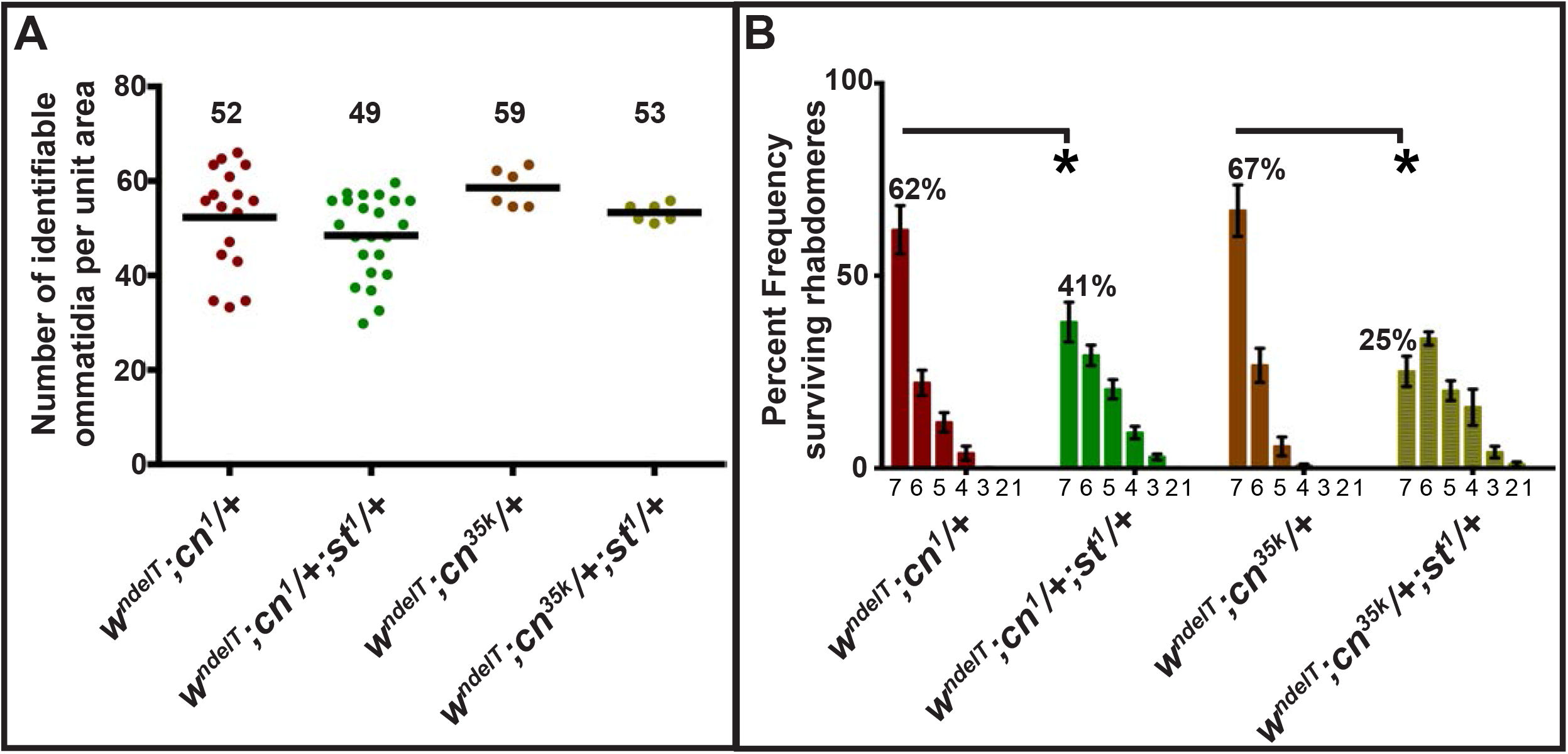
*cn^35K^* modulates retinal degeneration to the same extent as *cn^1^*. **A:** Graph comparing the extent of retinal damage following exposure to continuous, high intensity, white light for 7 days. Horizontal bars indicate the average number of ommatidia with identifiable rhabdomeres normalized to unit area. Each circle represents an individual count from 1 biological replicate. Number indicates the mean value for each condition. No statistically significant differences were recorded between *cn^1^/+* and *cn^35k^/+*. **B:** Percent frequency of ommatidia [± s.e.m.] displaying 1-7 rhabdomeres upon 7 days of continuous, high intensity, white light. Number on top of each bar indicates the mean value of ommatidia displaying the full complement of 7 rhabdomeres. Statistically significant differences, between genotype pairs as revealed by ANOVA followed by Bonferroni’s post-hoc test at p<0.05, are indicated by *. No statistically significant differences were recorded between *cn^1^/+* and *cn^35k^/+*.

**Supplemental Figure S3:**
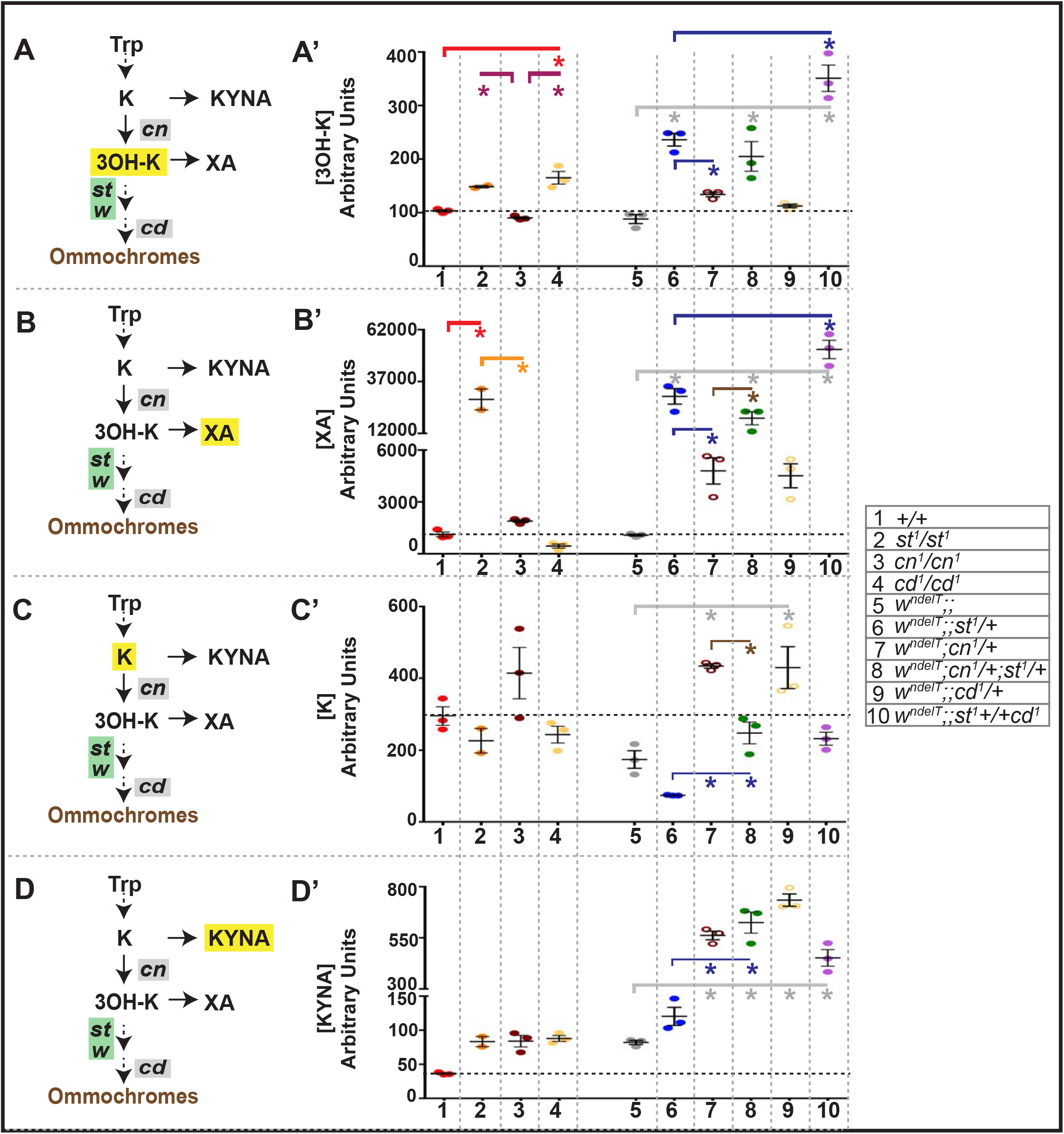
Mass Spectrometric (M.S.) measurements of select metabolites of the Kynurenine pathway (KP) **A, B, C, D:** Outline of relevant steps in the KP and brown pigment biosynthesis and genes analyzed in this study. The metabolite quantified and compared across genotypes in the corresponding graph (A’-D’) is highlighted in yellow. **A’-D’**: Graphs depicting the abundances of metabolites (highlighted in yellow in the corresponding pathway shown in A-D) from M.S. measurements (in arbitrary units along the Y axis) of head extracts. Numbers on X-axis correspond to the genotypes listed in the table on the right. Each colored dot represents an individual biological replicate of 10 pooled heads. Statistical comparisons between genotype pairs are indicated with solid lines (red line compared to +/+, magenta line (A’) compared to *cn^1^/cn^1^*, orange line (B’) compared to *st^1^/st^1^*, grey line compared to *w^ndelT^*, blue line compared to *w^ndelT^;;st^1^/+*, brown line compared to *w^ndelT^;cn^1^/+*). Statistically significant differences between pairs of genotypes are indicated by * as revealed by ANOVA followed by Bonferroni’s post-hoc test at p<0.05.

**Supplemental Figure S4:**
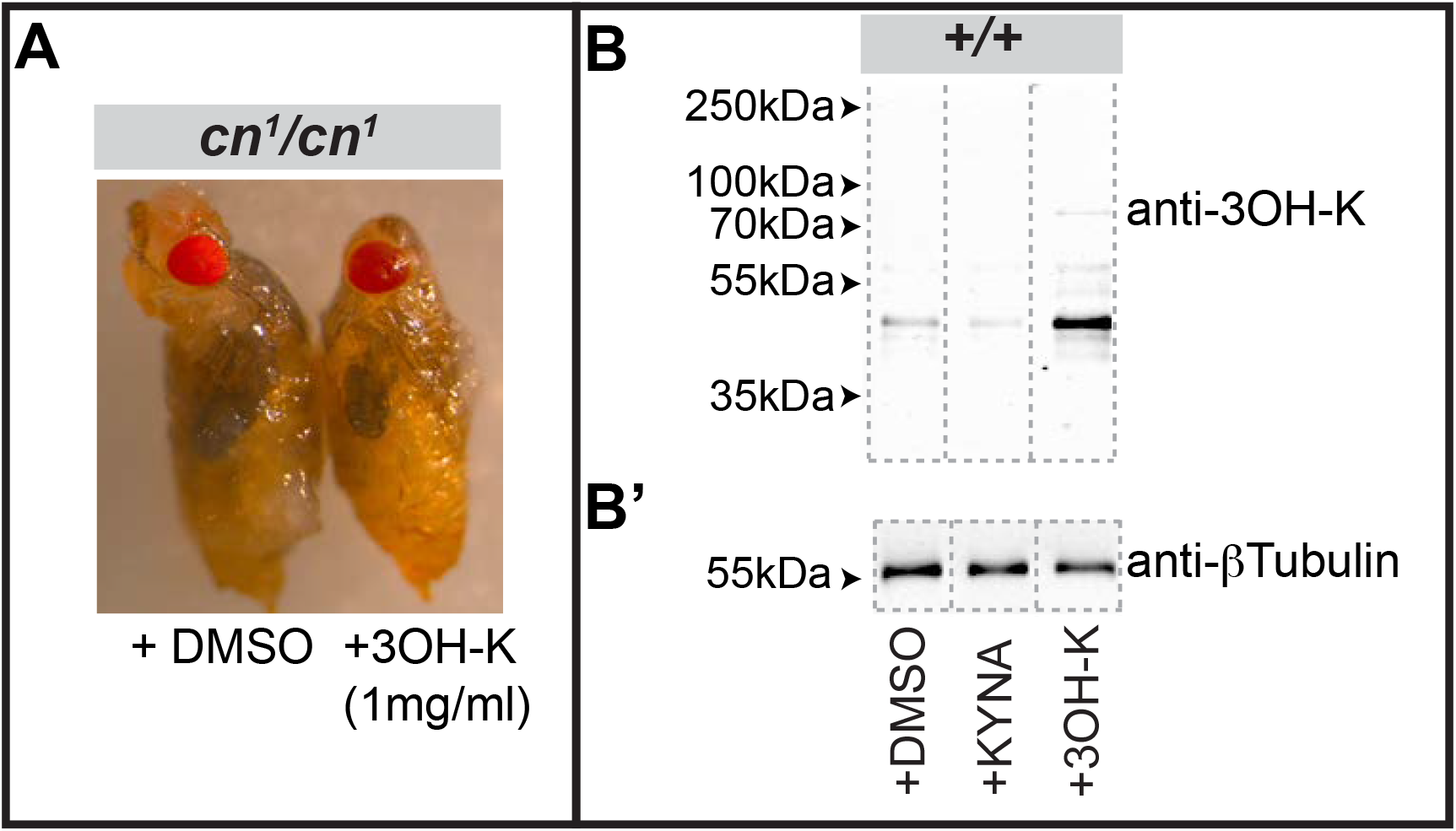
Effect of dietary supplementation of 3OH-K on eye color and the amount of protein-bound 3OH-K. **A:** Bright-field images of *cn^1^/cn^1^* flies raised on control food (+DMSO) or on food supplemented with 1mg/ml 3OH-K (+3OH-K). Control food-fed flies exhibit bright red eye color indicative of incomplete brown pigment biosynthesis (as expected of *cn* perturbation). Dietary 3OH-K reverts the eye color to dark red, indicative of proper brown pigment biogenesis. **B-B’:** Western blots of protein extracts from heads of wildtype adult flies (+/+) raised on control food (+DMSO), or on food supplemented with either 1mg/ml KYNA or 1mg/ml 3OH-K. Blots were tested with antibodies against 3OH-K (B) and against β-Tubulin as loading control (B’) run on parallel gels. Positions of standard molecular weight markers are indicated by arrowheads. Grey dotted lines outline the lanes of each sample.

**Supplemental Figure S5:**
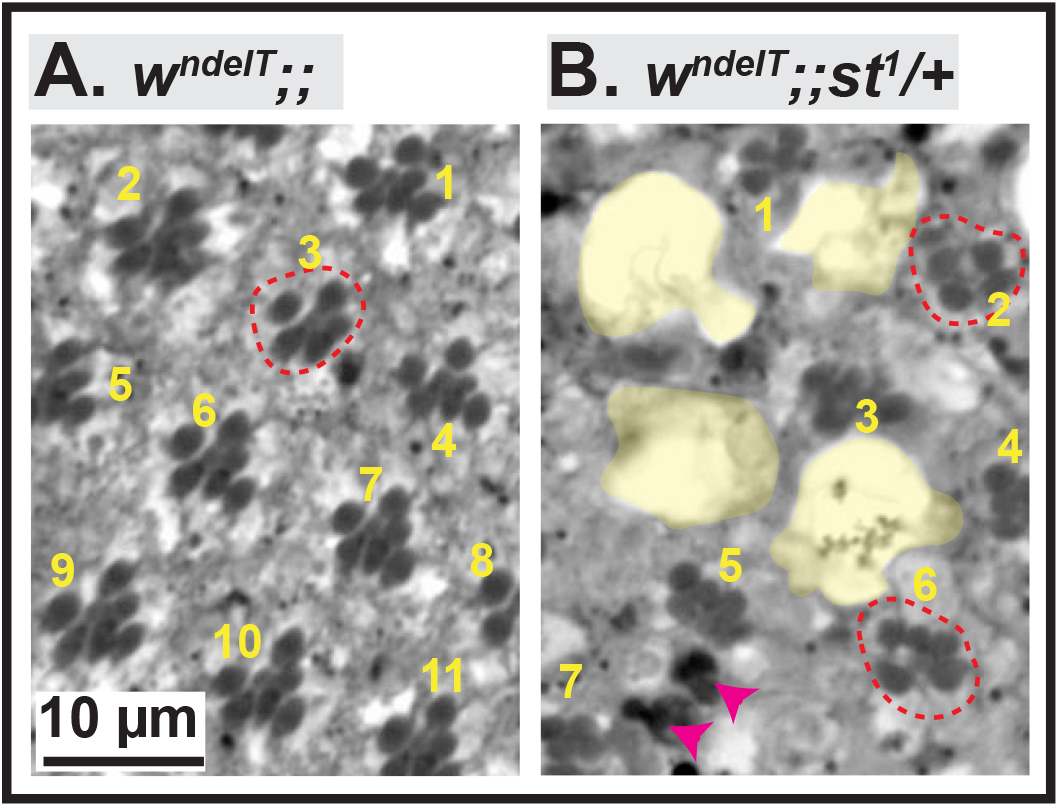
The influence of light exposure on retinal histology. **A-B:** are examples of images of toluidine-blue stained 1μm thin sections of fly eyes of *w^ndelT^* (A) and *w^ndelT^;;st^1^/+* (B), following exposure to high intensity continuous white light for 7 days. Qualitatively varying degrees of light induced damage are observed including holes (lacunae) (highlighted in yellow in B), ommatidia with less than 7 rhabdomeres (red dotted circles), and the presence of apoptotic debris (intensely stained regions; arrows in B). Scale bar as indicated in A. These phenotypes appear more pronounced in B than in A. Of these attributes, we quantified the following: (i) the consequence of lacunae formation by estimating ommatidial density or the number of identifiable ommatidia per unit area (11 in A, 7 in B). (ii) the status of photoreceptor (PRC) health by estimating the frequency of ommatidia with 7 or fewer rhabdomeres.

## Materials and Methods

### Fly strains and genetics

All phenotypic analyses were performed in age-matched (2 days of adulthood) males, unless otherwise specified, during the light-cycle of the incubator. Flies were maintained at 25°C on standard food (In water: corn flour-8%, malt-8%, sugar beet molasses-2.2%, soy-1%, yeast-1.8%, agar-0.75%, propionic acid-0.625%, 30% Nipagin (in ethanol)-0.5%). Wild-type flies were from the *Oregon R* strain. The *white* allele, designated as *w^ndelT^;; or w^ndelT^*, as it is a null allele that carries a deletion removing the transcriptional and translation start site of the *w* gene (Hebbar et al., 2021). Other mutant lines were obtained from The Bloomington *Drosophila* Stock Center and include: *cn^1^* (Line ID: BDSC_263), *cn^35k^* (Line ID: BDSC_268), *st^1^* (Line ID: BDSC_605) and *cd^1^* (Line ID: BDSC_3052). Double mutant heterozygous combinations were made by crossing female flies (*w^ndelT^*) with males of each of the mutant lines and selecting age-matched male flies. A homozygous line bearing *w^ndelT^* and *st^1^*mutations (*w^ndelT^;;st^1^*) was generated. Male flies of triple mutant combinations were generated by crossing female flies of *w^ndelT^;;st^1^* to male flies of either *cn^1^*, *cn^35k^, or cd^1^*. RNAi-mediated knockdown of the *white* gene was achieved with a line in which the GMR promoter drives the expression of white double stranded-hairpin RNA ((Lee & Carthew, 2003), The Bloomington *Drosophila* Stock Center Line ID: BDSC_32067, called *wRNAi* here). RNAi-mediated knockdown of the *st* gene was achieved by driving the expression of transgene bearing an inverted repeat of *st* (Cunningham et al., 2018, Dietzl, Chen et al., 2007) with the *GMR-Gal4* line (Freeman, 1996).

### 3OH-K and KYNA feeding

Animals were raised and maintained from embryonic stages onward on normal food supplemented with 3OH-K (1mg/ml of 3-Hydroxy-DL-kynurenine, H1771, Sigma Aldrich; dissolved in DMSO) or KYNA (1mg/ml of Kynurenic acid, K3375, Sigma Aldrich; dissolved in water).

### Experimental light conditions

Light conditions used here have been previously described (Hebbar et al., 2021). Briefly, flies were reared in regular light conditions defined as 12 hours of light (approx. 900-1300 lux)/12 hours of darkness. For the light-induced degeneration paradigm, flies (2-4 days of age) were placed at 25°C for 7 days in an incubator dedicated for continuous, high intensity light exposure (Johnson, Grawe et al., 2002b). High intensity light was defined by 1200-1300 lux measured using an Extech 403125 Light ProbeMeter (Extech Instruments, USA) with the detector placed immediately adjacent to the vial and facing the nearest light source. To remove blue light in this setting (Fig. 3), a customized box bounded by filters, which block blue light and face the light source in the incubator, was used. Light intensity was determined by measuring light counts using a USB spectrometer (Ocean Optics, USA).

### Histology and Quantification from toluidine blue-stained sections

Histology and analyses of toluidine blue stained sections were carried out as previously described (Hebbar et al., 2021) and modified from (Bulgakova, Rentsch et al., 2010). To compare morphological aspects of degeneration across different conditions, two aspects of light-induced damage were quantified. These include

1. the number of identifiable ommatidia normalized to area. Light exposure causes holes (highlighted in yellow in Fig. 2B) between adjacent ommatidia [also called lacunae (Ferreiro et al., 2017)] and it results in the reduced density of ommatidia. Therefore the quantification of the density of ommatidia is a measure of the consequence of holes in the retinal tissue. For example, within the same area, a relatively undamaged retina (*w^ndelT^*) exhibits 11 identifiable ommatidia (Suppl. Fig. S5) as opposed to only 7 identifiable ommatidia in a retina (*w^ndelT^;;st^1^/+*) with large holes (Suppl. Fig. S5) following light exposure. For the quantification, number of ommatidia were normalized to unit area x 10.
2. the number of rhabdomeres per ommatidia. The number of identifiable rhabdomeres per ommatidia was counted to calculate the percent frequency of ommatidia with fewer than the normal complement of 7 rhabdomeres. This attribute is recorded as a proxy of photoreceptor (PRC) health. For example, in wild-type retinas 100% of ommatidia showed seven rhabdomeres/ommatidium in each section (Fig. S1A’), indicative of seven healthy PRCs. It should be noted that fewer than 7 rhabdomeres per ommatidia does not necessarily indicate cell death.

### Metabolite extraction, MS measurement

10 heads from age-matched (2-4 days old) male flies of genotypes indicated were pooled as one biological replicate and three such replicates were generated. Fly heads were frozen in liquid nitrogen and kept at −80°C until further use. Fly heads were homogenized using 0.5 mm zirconia beads for 2 x 5min in 100μl 80% methanol; 0.5% Formic acid and 50nM Internal standard Chloropropamide (Sigma-Aldrich Chemie GmbH). The mixture was centrifuged at 13.000g, and 10μl were analyzed. Mass spectrometric analysis was performed on a Q Exactive instrument (Thermo Fischer Scientific, Bremen, DE) equipped with a robotic nanoflow ion source TriVersa NanoMate (Advion BioSciences, Ithaca, USA) using nanoelectrospray chips with a diameter of 4.1 μm. The ion source was controlled by the Chipsoft 8.3.1 software (Advion BioSciences). Ionization voltage was + 0.96 kV in positive and - 0.96 kV in negative mode; backpressure was set at 1.25 psi in both modes. Samples were analyzed by polarity switching (Schuhmann, Almeida et al., 2012). The temperature of the ion transfer capillary was 200 °C; S-lens RF level was set to 50%. All samples were analyzed for 11 min. FT MS spectra were acquired within the range of m/z 150–350 from 0 min to 0.2 min in positive and within the range of m/z 150–350 from 6.2 min to 6.4 min in negative mode at the mass resolution of R m/z 200 = 140000; automated gain control (AGC) of 3×106 and with the maximal injection time of 3000 ms. t-SIM in positive (0.2 to 4.2 min) and negative (6.4 to 10.4 min) mode was acquired with R m/z 200 = 140000; automated gain control of 5 × 104; maximum injection time of 650 ms; isolation window of 20 Th and scan range of m/z 150 to 350 in positive and m/z 150 to 350 in negative mode, respectively. The inclusion list of masses targeted in t-SIM analyses started at m/z 150 in negative and m/z 150 in positive ion mode and other masses were computed by adding 10 Th increment (i.e. m/z 155, 165, 175, …) up to m/z 350. Quantification of all targeted molecules was performed by parallel reaction monitoring (PRM) FT MS/MS from 4.2 to 6 min in positive mode and from 10.4 to 11.0 min in negative mode. For PRM micro scans were set to 1, isolation window to 0.8 Da, normalized collision energy to 12.5%, AGC to 5 × 104 and maximum injection time to 650 ms. All spectra were pre-processed using repetition rate filtering software PeakStrainer (Schuhmann, Thomas et al., 2017) and stitched together by an in-house developed script (Schuhmann, Srzentic et al., 2017). The targeted molecules were quantified by LipidXplorer software (Herzog, Schuhmann et al., 2012).

### Western Blotting

Three fly heads from age-matched flies were homogenized in 10 μl of 4x SDS-PAGE sample buffer (200 mM Tris-HCl pH 6.8, 20% Glycerol, 8% SDS, 0.04% Bromophenol blue, 400 mM DTT). After dilution with RIPA buffer (150 mM sodium chloride, 1% Triton X-100, 0.5% sodium deoxycholate, 0.1% SDS, 50 mM Tris pH 8), lysates were heated at 37°C for 30 min. Lysates equivalent to 2.5 heads were loaded and run on two parallel (15% acrylamide) gels, and proteins were transferred onto a membrane (Nitrocellulose Blotting Membrane 10600002; GE Healthcare Life Sciences; PA; USA). Primary antibodies were incubated overnight at 4°C and included mouse anti-3OH-K (1:1000; MA1-16616, AB_568420, clone P3UI, Thermo Fisher Scientific) and mouse anti-β-Tubulin E7 (1:1000; [http://dshb.biology.uiowa.edu/tubulin-beta-_2] deposited by M. McCutcheon/S. Carroll to the from Developmental Studies Hybridoma Bank (DSHB), University of Iowa, USA). As secondary antibody IRDye 800CW goat anti-Mouse IgG (1:15,000; LI-COR Biotechnology; NE; US) was used for an 1 h incubation at room temperature. The dry membrane was imaged using a LI-COR Odyssey Sa Infrared Imaging System 9260-11P (LI-COR Biotechnology).

### Figure panel preparation and statistical analyses

Statistical power analyses was not conducted as part of experimental design to determine sample size. Sample sizes are evident in all dot plot graphs. For frequency distribution, sample sizes are indicated in the legend. All figure panels were assembled using Adobe Photoshop CS5.1 or Adobe Illustrator CS3 (Adobe Systems, USA). Statistical analyses and graphs were generated using GraphPad Prism (GraphPad Software, Inc, USA).

## Notes

The authors declare that they have no conflict of interest.

